# Gradual evolution of multiplexed spatial coding within a conserved hippocampal scaffold

**DOI:** 10.64898/2026.09.19.752914

**Authors:** Xiao-Fan Ge, Chu Deng, Jia-Wei Zou, Yi-Qing Hui, Yi-Fan Ye, Jia-Li Long, Qiu-Chi Xu, Ya-Li Duan, Shu-Yi Hu, Wei-Bo Kang, Wen-Cong Li, Kun Jiang, Ci-Rong Liu, Yong-Gang Yao, Xing Cai, Bai-Lu Si, Li Lu

## Abstract

The hippocampus constructs spatial maps for navigation, yet their format differs markedly across species. Whether divergences reflect ecological adaptation, phylogenetic architecture, or flexible coding strategy within a conserved circuit has been difficult to disentangle. We recorded CA1 in rats and tree shrews (*Tupaia belangeri chinensis*), a mammal phylogenetically and ecologically between rodents and primates, using identical tasks and analyses. Unlike the position-dominant code of rats, tree shrew CA1 exhibited a hybrid representation: weaker position selectivity, stronger non-positional tuning, and prevalent multiplexing approaching primate characteristics. Multiplexed populations decoded location as accurately as rat position populations while using fewer neurons, suggesting computational advantages of high-dimensional coding. Despite this representational shift, proximo-distal gradient, pattern completion, and global remapping persisted. Representational format can be tuned without altering the underlying circuit. Tree shrews occupy an evolutionary intermediate in which ancestral network dynamics are repurposed for efficient multiplexed coding, tracing a transition toward primate-like spatial representations.

## INTRODUCTION

The hippocampus constructs internal spatial maps essential for navigation across mammals ^1, 2^, yet the format of these maps differs dramatically between species. Rodent hippocampal pyramidal neurons primarily encode position via discrete place fields ^3^, whereas monkey hippocampal neurons represent multiple spatial variables including position, head direction (HD), facing location (FL), spatial view (SV), and self-referenced motion cues ^4–9^. Human intracranial recordings have similarly revealed place, HD, and SV cells in the hippocampal formation ^10–12^. These species differ in both phylogeny and sensory ecology: rodents are largely nocturnal and rely on tactile and olfactory cues ^13^, whereas primates are mostly diurnal and visually guided ^6^. However, even comparisons involving freely navigating animals have employed divergent environments and tasks: rodents typically explore small, two-dimensional enclosures via self-locomotion ^14–16^, whereas freely navigating monkeys have been tested in larger, more complex layouts with distinct sensory and motor requirements ^5, 6, 9, 17^. These confounds have made it difficult to disentangle intrinsic neural organization from extrinsic behavioral demands.

Despite these methodological variations, two consistent distinctions emerge. First, rodent hippocampal principal neurons predominantly encode immediate position in physical space ^18, 19^, whereas primate representations exhibit weaker position modulation but stronger engagement with extrapersonal visual space ^20–22^. Second, primate hippocampal neurons frequently exhibit multiplexed tuning, with single cells simultaneously representing multiple spatial parameters ^5–8^, whereas most rodent hippocampal neurons appear predominantly position-modulated ^23, 24^. Whether these divergences reflect ecological pressures, phylogenetic differences in neural architecture, or flexible optimization of coding strategy within a conserved circuit remains unknown, as does whether the transition from rodent-like to primate-like coding occurs gradually or abruptly.

To address both the mechanism and pace of this transition, we compared hippocampal CA1 activity in Long-Evans rats and Chinese tree shrews (*Tupaia belangeri chinensis*). Tree shrews sit between rodents and primates both phylogenetically ^25, 26^ and ecologically: as rat-sized, semi-arboreal, diurnal mammals, they are intermediate between nocturnal rodents and visually guided primates ^13, 27^. This dual intermediacy makes tree shrews uniquely suited to reveal whether intermediate spatial coding exists, although a single species cannot separate the contributions of ancestry and ecology, a dissociation that will require comparing additional species. Using identical tasks, recording methods, and analytical tools in matching environments, we asked whether tree shrews exhibit hybrid representations bridging rodent and primate extremes. If multiplexed coding reflects a flexible adjustment of spatial representations within an evolutionarily conserved circuit, rather than a primate-specific innovation, tree shrews should show intermediate multiplexing alongside otherwise preserved network dynamics. Such a result would mean that richer spatial codes can arise without wholesale changes to circuit architecture, separating representational format from the computations that generate it, and would help explain how the hippocampus adapts to diverse ecological demands while retaining its core spatial function.

## RESULTS

### Comparable recording sites between tree shrews and rats

We employed a custom-designed tetrode drive with wireless logging to monitor dorsal CA1 activity in freely behaving tree shrews (Fig. 1a). Both species were tested in a standard 1 m × 1 m open arena with a proximal cue card and distal cues obscured by curtains (Fig. 1b). We recorded from 14 young adult male tree shrews (25.1 ± 1.9 months old) and 20 young adult male Long-Evans rats (26.9 ± 1.1 weeks old). Rat data were reported previously ^19^ but are re-analyzed here using different methods.

**Fig. 1.**
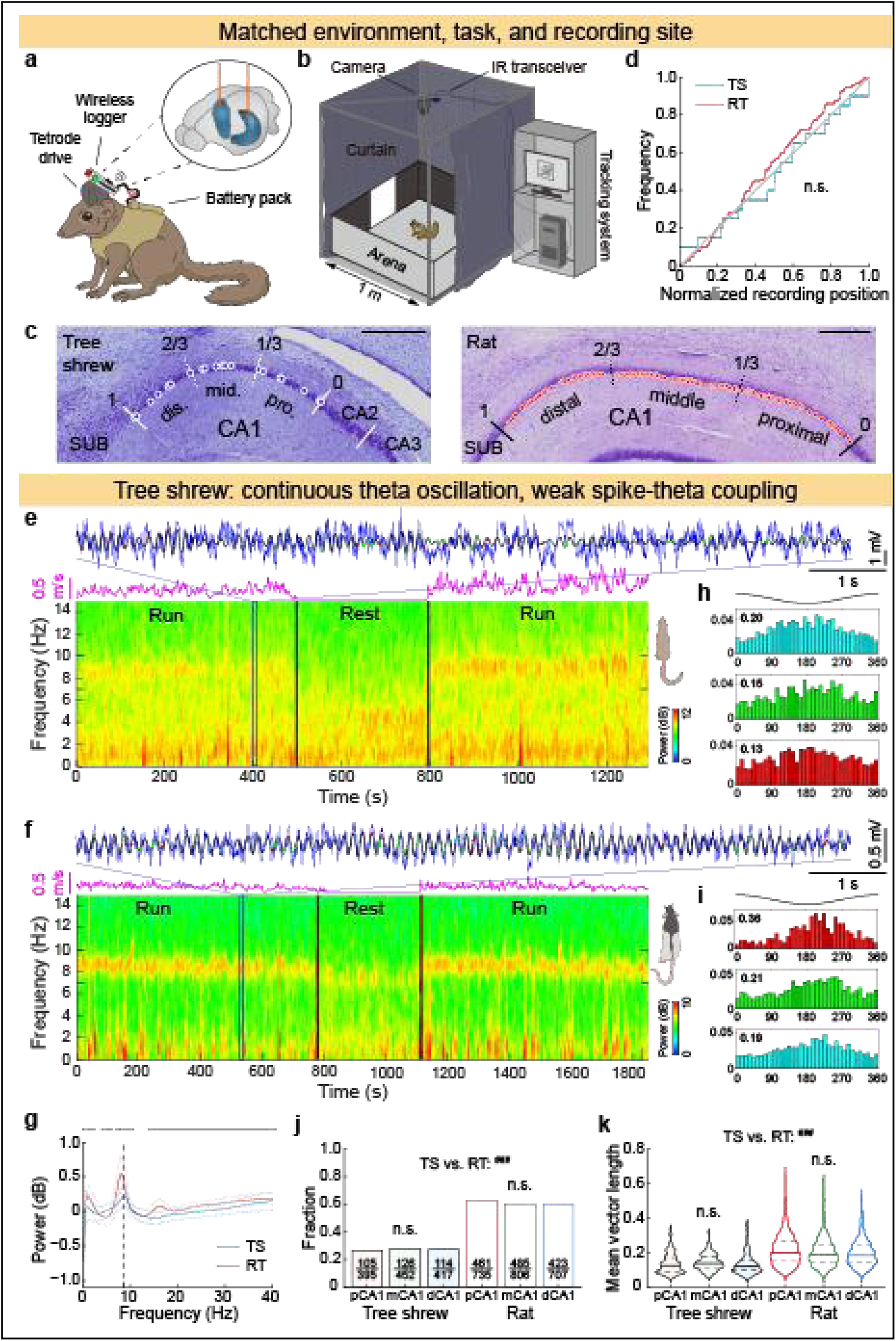
Continuous theta with attenuated synchrony in tree shrew CA1. **a,** Wireless tetrode drive for freely behaving tree shrews. **b**, Standard 1 m^2^ open arena with proximal cue card and curtained distal cues. **c**, CA1 transverse axis recording sites. Dots, tetrode tips; numbers, normalized positions. SUB, subiculum. Scale: 0.5 mm. Detailed rat histology: see ^19^. **d**, Cumulative site distributions. n.s., not significant. **e**–**f**, Representative theta for tree shrews and rats. Top: 10-s raw local field potential (LFP; blue) and theta-band (7–10 Hz) filtered signal (black) recorded during active navigation. Colored dots, spike times of neurons in **h**, **i**. Middle: Instantaneous speed (magenta). Bottom: spectrogram during locomotion and rest (dashed: 7–10 Hz). **g**, Aperiodic-removed power spectra (mean ± SD). Dashed line: 8.5 Hz. Gray bars: significant between-species differences. **h**–**i**, Example theta phase-locked neurons (mean vector length (MVL) in bold), with preferred firing at troughs. Abscissa: theta phase; ordinate: frequency. **j**, Proportion of theta phase-locked neurons. RT, rat; TS, tree shrew. **k**, Theta phase-locking strength (MVL; median (IQR)). Tree shrews exhibit rodent-like continuous theta but primate-like weak spike-theta coupling. n.s., not significant; cross-region: \**P* < 0.05, \*\*\**P* < 0.001; cross-species: ^###^*P* < 0.001.

Recording sites spanned the CA1 transverse axis in both species (Fig. 1c, Extended Data Fig. 1) and were distributed uniformly (Fig. 1d; Supplementary Table 1). We normalized positions relative to the CA1/CA2 border and classified recordings as proximal (pCA1, 0–0.333), middle (mCA1, 0.333–0.667), or distal (dCA1, 0.667–1) ^16^. In tree shrews, we obtained 422, 483, and 420 active (mean rate ≥ 0.1 Hz) excitatory neurons from 7 pCA1, 6 mCA1, and 7 dCA1 sites, respectively. Rat datasets included 863, 940, and 773 neurons from 39 pCA1, 43 mCA1, and 31 dCA1 sites, using the same clustering method (Extended Data Fig. 2). This balanced sampling rules out regional bias.

### Continuous theta with attenuated spike-field synchrony in tree shrew CA1

Rodents exhibit prominent, continuous theta (7–10 Hz) during navigation, whereas primate theta occurs in brief, intermittent, movement-modulated bouts ^5, 28–30^. Tree shrews displayed robust, continuous theta oscillations peaking at ∼8.5 Hz during locomotion, markedly attenuated during rest (Fig. 1e), qualitatively resembling rat theta dynamics (Fig. 1f) but with substantially lower relative power after removing aperiodic components ^31^ (Fig. 1g).

A substantial proportion of tree shrew CA1 principal neurons fired preferentially at the theta trough (Fig. 1h), as in rats (Fig. 1i) and macaques ^5^. In line with lower theta power, the fraction of neurons exhibiting significant theta phase-locking (Rayleigh test, *P* < 0.01) was substantially lower in tree shrews than in rats (27.3% vs. 60.9%; Fig. 1j), comparable to macaques (∼25%)^5^. The strength of theta phase-locking (mean vector length, MVL) was also markedly weaker in tree shrews, with no clear proximo-distal gradient in either species (Fig. 1k). Thus, tree shrew hippocampus generates rodent-like theta oscillations but with primate-like weak spike-theta synchrony.

### Distinct navigation strategies in tree shrews and rats

Both species explored the arena freely using quadrupedal locomotion to forage for small food crumbs, yet displayed distinct strategies. Tree shrews maintained forward gaze and traversed the arena in relatively straight paths under normal illumination (Supplementary Video 1), whereas rats exhibited rhythmic lateral head sweeping to scan the environment with their whiskers and followed meandering trajectories under dim light (Fig. 2a). Therefore, tree shrews showed higher linear speed (LS) and angular velocity (AV) (Fig. 2b–d; Supplementary Table 1), whereas rats displayed greater AV/LS ratio, path tortuosity, and thigmotaxis (Fig. 2e–g). Neither species exhibited prolonged stationary periods with saccadic head scanning characteristic of primate visual exploration ^6^ (Fig. 2g). Despite straighter, faster trajectories, tree shrews sampled the arena as uniformly as rats (Fig. 2h, i), ruling out differential coverage as an explanation for coding differences.

**Fig. 2.**
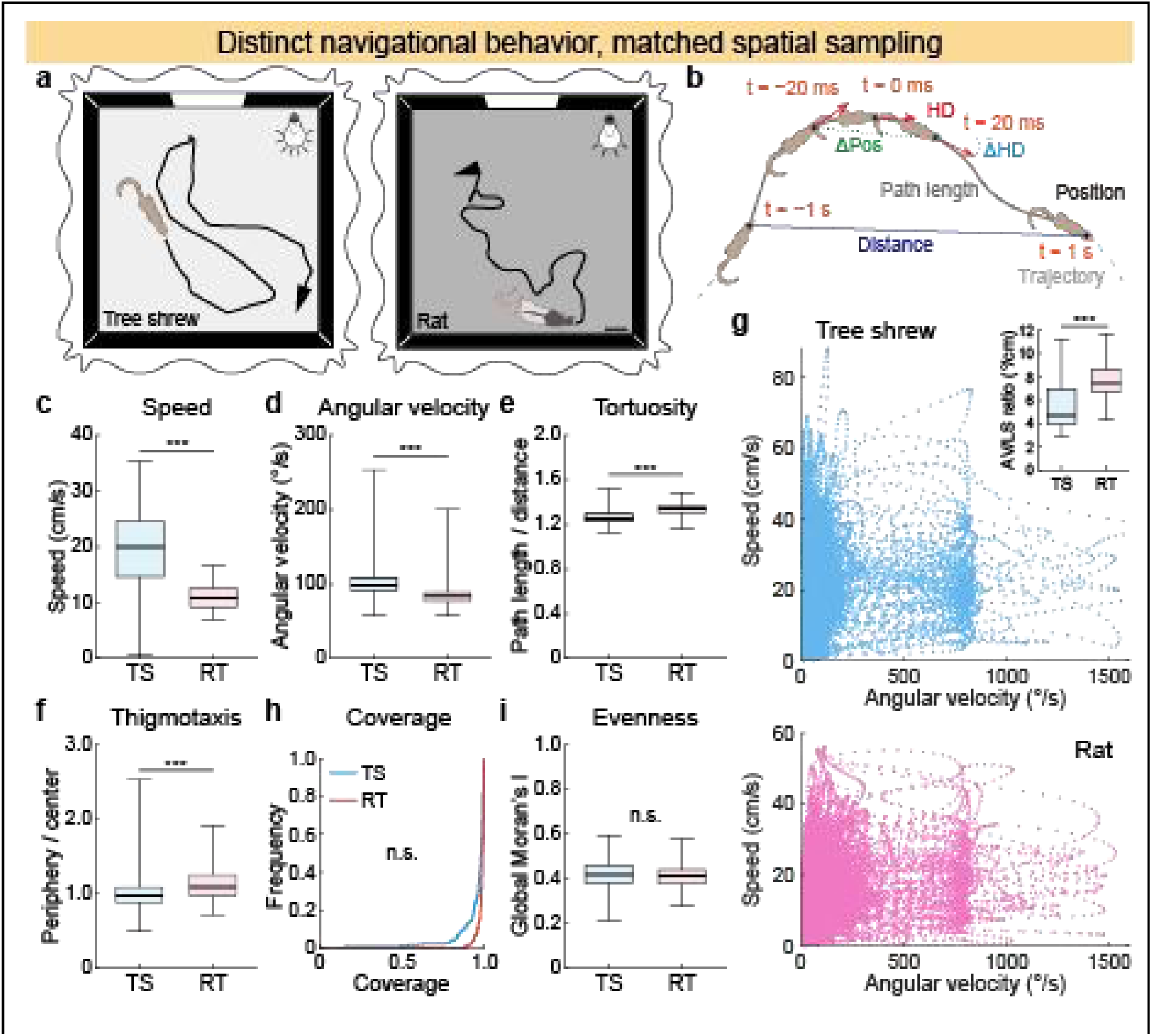
Distinct navigation in tree shrews and rats. **a**, Representative 10-s trajectories for both species. Scale bar: 10 cm. **b**, Behavioral metrics. Pos, position; HD, head direction. **c**–**f**, Species differences in linear speed (LS), angular velocity (AV), path tortuosity, and thigmotaxis (median (IQR)). **g**, Representative AV vs. LS scatters; neither species shows primate-like head scanning (high AV at low LS). Inset: AV/LS ratio. **h**–**i**, Comparable spatial coverage and evenness. Tree shrews ran faster and straighter; rats swept their heads and followed walls. Both species sampled the arena uniformly. n.s., not significant; \*\*\**P* < 0.001.

### Moderate, graded spatial modulation in tree shrew CA1

We next quantified basic firing properties and position selectivity of CA1 pyramidal neurons during navigation (spatial variables illustrated in Fig.3a). Mean firing rates were comparable between species, except in tree shrew dCA1, which may include subicular neurons from one tetrode at the CA1-subiculum border (Extended Data Fig. 3a). We identified tree shrew neurons that fired at specific positions in the arena, closely resembling classic rodent place cells (Fig. 3b). Notably, proximal neurons displayed single, coherent receptive fields, whereas distal neurons showed irregular, spatially dispersed activity. A similar proximo-distal gradient was present in rat CA1 ^16^, though rat neurons displayed stronger overall spatial selectivity, as evidenced by lower total spatial coverage per neuron (Extended Data Fig. 3b). Despite this shared decline, dCA1 field structure diverged: rat neurons tended to exhibit multiple discrete fields, whereas tree shrew neurons typically showed a single extended field (Extended Data Fig. 3c).

**Fig. 3.**
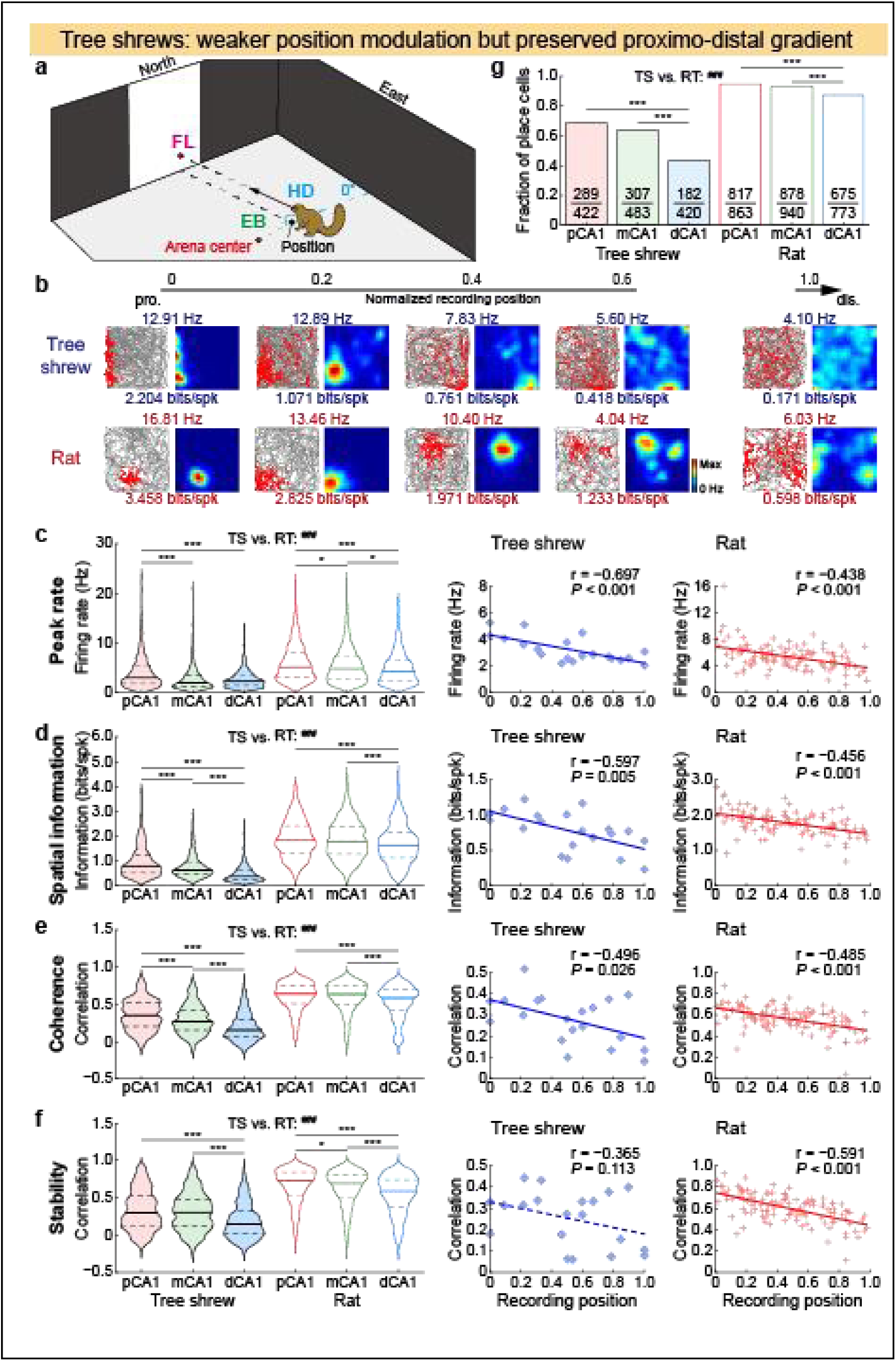
Moderate, graded spatial modulation in tree shrew CA1. **a**, Spatial variables: position, HD, facing location (FL), and egocentric bearing (EB). **b**, Example cells showing spike-trajectory plots (left; path: gray; spikes: red) and color-coded rate maps (right). Top numbers, peak firing rates; bottom, spatial information content (SIc). **c**–**f**, Position selectivity metrics. Left: violin plots by subregion (median (IQR)). Right: metrics vs. normalized recording positions; solid lines, significant regression; dashed, non-significant. No species difference in transverse slope. **g**, Fractions of position-modulated neurons by subregion (numbers, neuron counts). Tree shrew CA1 shows weaker but graded position tuning with conserved proximo-distal organization. Cross-region: \**P* < 0.05, \*\*\**P* < 0.001; cross-species: ^###^*P* < 0.001.

We compared spatial modulation using peak firing rates (Fig. 3c), spatial information content (SIc; Fig. 3d), spatial information rate (Extended Data Fig. 3d), spatial coherence (Fig. 3e), and spatial stability (Fig. 3f). All measures were significantly lower in tree shrews (left panels, two-way rank-transformed analysis of variance (RT-ANOVA); species effect: all *P* < 0.001), confirming reduced position selectivity. Both species exhibited gradual decreases from pCA1 to dCA1 (subregion effect: all P < 0.001), and the magnitude of these gradients differed between species for most metrics (species × subregion interaction: all *P* ≤ 0.004, except spatial stability; Supplementary Table 1). Similar results were obtained using linear mixed-effects (LME) models, although pCA1-to-dCA1 differences were attenuated (Supplementary Table 2), suggesting that inter-individual variation in recording site distribution contributed to gradient estimates. The proximo-distal differences were also evident at the tetrode level: averaging across neurons at each recording site revealed consistently weaker spatial modulation in tree shrews and comparable transverse gradients between species (permutation test; all *P* > 0.1; Fig. 3c–f & Extended Data Fig. 3d, right panels).

We next quantified the proportions of position-modulated neurons using a permutation test based on SIc. The proportion of position-modulated neurons in tree shrew CA1 (43.3–68.5%) was lower than in our rat recordings (87.3–94.7%) and in previously reported mouse CA1 (> 90%) ^18^, but higher than the position-coding fractions reported in freely moving monkeys (14–r32%) ^5, 6, 9, 17, 29, 32^. Both species showed higher place cell fractions in pCA1 than in dCA1 (Fig. 3g). The moderate position tuning in tree shrews was not attributable solely to fewer place cells; position modulation strength was also significantly weaker among neurons that met classification criteria (Extended Data Fig. 4a–h).

Together, tree shrew CA1 exhibits moderate position selectivity, consistent with a graded evolutionary decline in position tuning from rodents to primates. At the same time, the preserved proximo-distal gradient indicates that the core functional organization of the hippocampus remains conserved despite divergent spatial codes.

### Enhanced non-positional spatial coding in tree shrew CA1

Unlike rodents, primate hippocampal neurons encode multiple non-positional variables, including HD, FL (approximating SV), and egocentric boundaries during navigation ^5–9^. We systematically examined these variables in tree shrews (Fig. 3a). Among position-modulated neurons, a substantial proportion also responded to HD, FL, and/or egocentric bearing (EB) relative to the arena center. By contrast, rat place cells typically encode position predominantly, with minimal conjunctive coding (Fig. 4a).

**Fig. 4.**
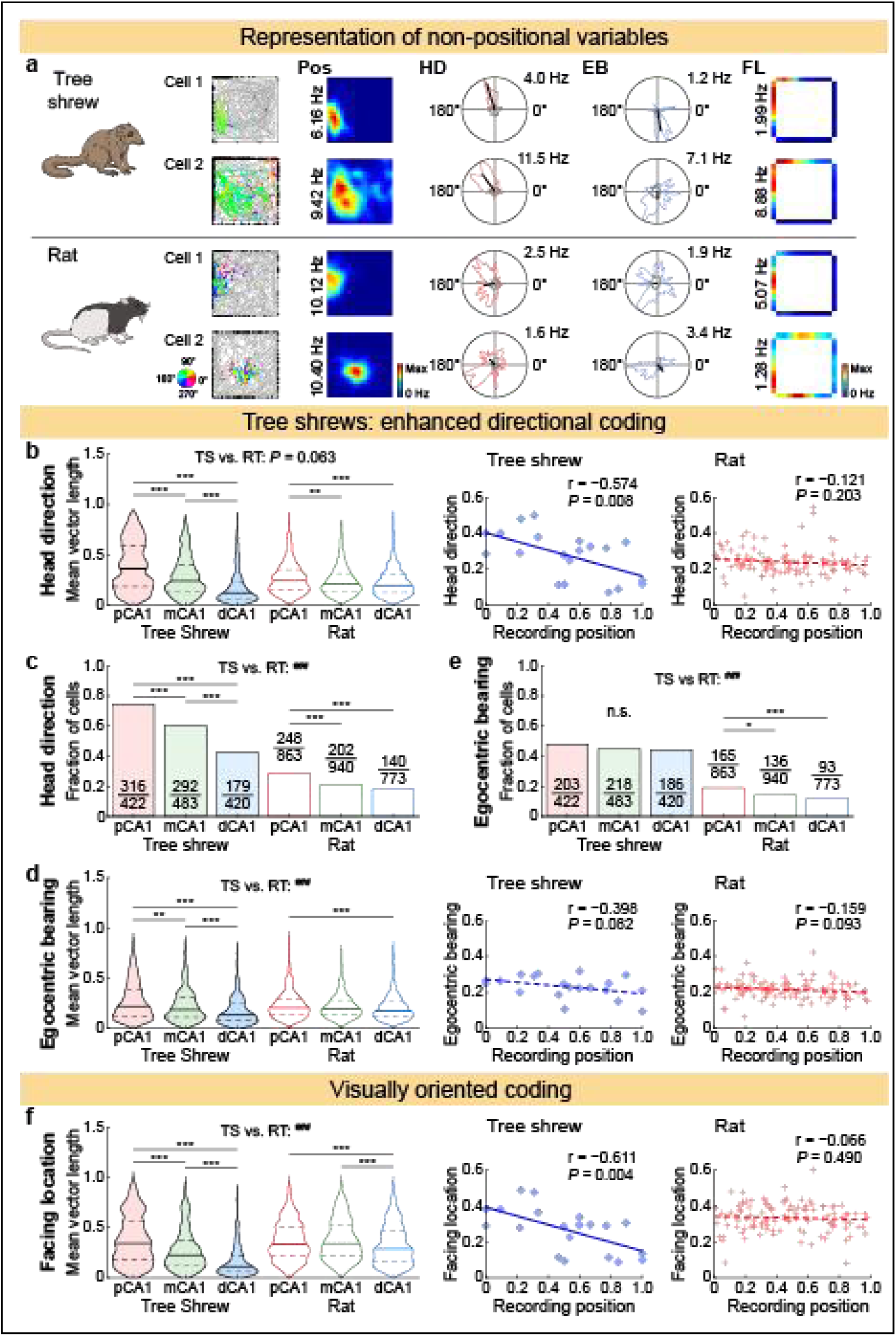
Graded non-positional coding along the CA1 proximo-distal axis. **a**, Representative neurons: tree shrews show multiplexed non-positional tuning; rats show predominantly position tuning. Columns: (1) spike-trajectory plots (gray: path; colored dots: spikes with color-coded HD; black dots: FL projections); (2) position rate maps; (3) HD tuning curves (red; gray: time distribution; black line: MVL and preferred HD); (4) EB tuning curves (blue); (5) FL rate maps. Peak rates indicated. **b**–**c**, HD tuning strength (MVL) and proportion of HD-tuned neurons. **d**–**e**, EB tuning strength and proportions. **f**, FL tuning strength. Tree shrews contain more HD- and EB-modulated neurons than rats, with steeper proximo-distal gradients. Cross-region: \*\**P* < 0.01, \*\*\**P* < 0.001; cross-species: ^###^*P* < 0.001.

We computed the MVL for each neuron’s tuning curve. This revealed substantially stronger HD tuning in tree shrew pCA1, declining markedly toward dCA1 (two-way RT-ANOVA; subregion: *P* < 0.001). Rats showed only moderate changes in HD modulation along the proximo-distal axis (species × subregion: *P* < 0.001; Fig. 4b, left; Supplementary Table 1). At the tetrode level, HD modulation changed more steeply across subregions in tree shrews, consistent with sharper functional segregation (Fig. 4b, right; permutation test, *P* < 0.01). Tree shrews contained significantly higher proportions of HD-tuned neurons (42.6–74.9%; identified via permutation testing) than rats (18.1–28.7%) (Fig. 4c). Unlike position coding, the modulation strength of HD-tuned neurons was comparable across species (Extended Data Fig. 4i).

Unlike allocentric HD coding, EB modulation was weaker in both species (spatial metric: *P* < 0.001) and varied modestly along the proximo-distal axis at the cellular level (subregion: *P* < 0.001), with the same trend at the tetrode level (*P* < 0.1 in both species; Fig. 4d). Tree shrews exhibited significantly higher proportions of EB-modulated neurons (44.3–48.1%) than rats (12.0–19.1%) (Fig. 4e), with comparable MVLs between rat CA1 and tree shrew pCA1 (Extended Data Fig. 4j). The MVL–proportion discrepancy in both HD and EB may reflect rhythmic lateral head sweeping in rats (Fig. 2), which could elevate directional MVLs without producing consistent, permutation-significant tuning. Tree shrew CA1 neurons thus encoded primate-like directional variables with higher prevalence.

Tree shrews showed FL-modulation patterns similar to HD coding, with graded tuning strength and significant proportions of modulated neurons along the CA1 transverse axis (Fig. 4f, Extended Data Fig. 4k, l). However, FL tuning can be confounded by place field proximity to boundaries, an effect more pronounced in rats, which display smaller, peripherally biased fields^14^ (Fig. 4a, cell 1). Accordingly, we assessed FL modulation primarily from generalized additive model (GAM) analyses that control for position.

### Multiplexed spatial coding in tree shrew CA1 but not rat CA1

Traditional tuning curve analysis assesses only one variable at a time, precluding rigorous assessment of multiplexed encoding in individual neurons ^24, 33^. We therefore adopted an unbiased, model-based framework using GAMs ^5, 24, 33, 34^, which evaluate multiple spatial variables concurrently while controlling for correlated artifacts.

The full model incorporated six variables: position, HD, FL, EB, LS, and AV (Figs. 2b, 3a). This model provided good fit (Extended Data Fig. 5a) and was more stringent than single-variable tuning curves, which credit neurons for apparent tuning that covaries with real tuning; accordingly, GAMs yielded lower proportions of spatially modulated neurons (Fig. 4c, e). Encoding was strongest in pCA1 in both species, and position remained the most robust variable. Within this shared framework, however, striking species differences emerged in complete encoding profiles. Tree shrew CA1 showed rich multiplexed coding of spatial variables, whereas rat CA1 remained predominantly position-focused (Fig. 5a, bar plots). Egocentric variable coding (EB, LS, AV) was minimal in both species (< 8%), indicating predominant allocentric coding by pyramidal neurons ^8^, consistent with interneuronal encoding of LS and AV ^6, 7, 29, 35, 36^. These findings were robust across classification criteria, including log-likelihood (goodness of fit) comparisons of first-order models (Extended Data Fig. 5b–e), and dominant variable fractions (Fig. 5a, pie charts; subregional data in Extended Data Fig. 6).

**Fig. 5.**
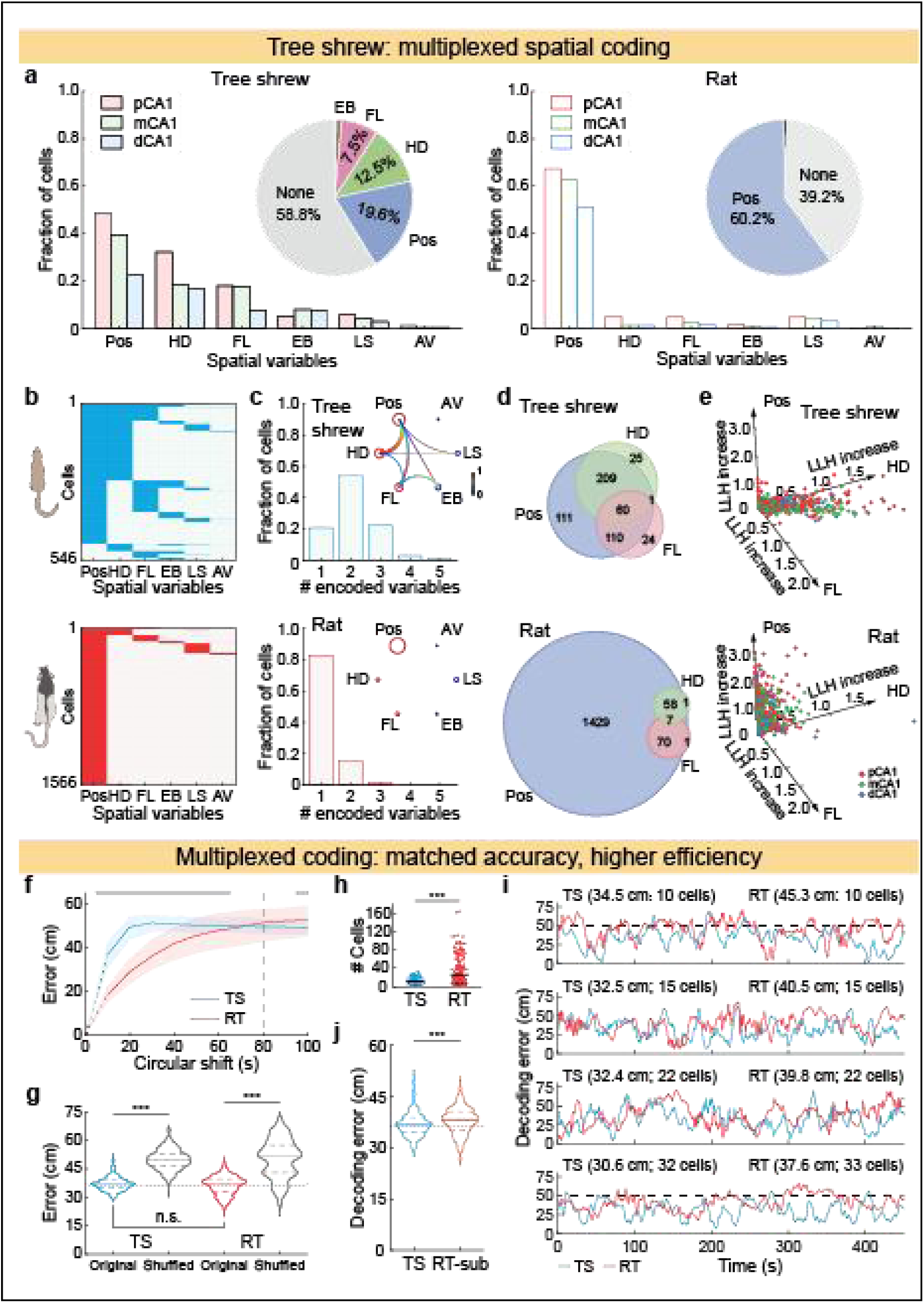
Multiplexed spatial coding in tree shrew but not rat CA1. **a**, generalized additive model (GAM)-derived proportions by complete encoding profile (bar plots) and dominant variable (pie charts). **b**–**d**, Multiplexed coding in tree shrews (top): spatial variable combination matrix, variable-count distribution, conjunctive coding circular graph and Venn diagram of allocentric tuning overlap (numbers, neuron counts). Bottom: corresponding analyses for rats. **e**, 3D scatter of Pos, HD, and FL log-likelihoods (values < 0 set to zero for better visualization). **f**, Temporal structure of navigational behavior. Error between original and time-shifted position trajectories as a function of circular time shift. Dashed line: 80 s; gray bars: significant between-species differences. **g**, Multi-session position decoding accuracy for the entire dataset (median (IQR)). Original vs. shuffled (spike-position circularly shifted by 80 s). **h**, Neurons per decoding session. TS, 8 (5–18) neurons; RT, 11.5 (7–38) neurons. **i**, Representative count-matched decoding error; individual tree shrews outperformed rats. **j**, Population count-matched decoding error across multi-session data. Multiplexed tree shrew populations achieve rat-comparable spatial accuracy with fewer neurons. n.s., not significant; \*\*\**P* < 0.001.

Among classified tree shrew spatial neurons, the majority were position-modulated, yet most additionally exhibited tuning to HD and/or other variables (Fig. 5b). Three-quarters were tuned to multiple variables: 20% encoding three variables and 55% encoding two (Fig. 5c, bar plot). The three allocentric variables (position, HD, and FL) were frequently co-coded within single neurons (Fig. 5c, circular graph). Among position-modulated neurons, 77.3% were co-coded with HD and/or FL. Allocentric co-coding was even higher among HD-modulated (91.5%) and FL-modulated (87.7%) neurons (Fig. 5d). In marked contrast, > 80% of rat spatial neurons were tuned exclusively to position, with sparse co-coding (Fig. 5b–d, bottom). GAM analyses thus revealed rich multiplexed representations of allocentric variables in tree shrew CA1, resembling primate hippocampal coding ^5–8^, whereas rat CA1 coding was almost exclusively position-based ^18, 23^ (Fig. 5e).

### Efficient spatial decoding by multiplexed representations

We next asked whether tree shrews benefited from multiplexed coding during navigation. Because wireless tree shrew recordings yielded smaller within-session populations than classical Bayesian decoders require, we adopted a multi-session data-sharing framework ^37^. To assess the temporal structure of navigational behavior, position trajectories were circularly time-shifted. Error plateaued at 20 s in tree shrews versus 80 s in rats (Fig. 5f), indicating faster position decorrelation during tree shrew navigation (Fig. 2). Despite this less predictable behavior, and thus a harder decoding problem, multiplexed tree shrew CA1 populations predicted instantaneous location as accurately as rat position-dominant populations across the full dataset, which was already matched for proximo-distal cell distributions between species (Fig. 5G; Supplementary Table 1), while doing so with fewer simultaneously recorded neurons per session (Fig. 5h). In a neuron count-matched comparison, rat decoding error remained significantly higher than that of tree shrews (Fig. 5i, j). Taken together, multiplexed coding achieves equivalent localization with sparser neural resources than position-dominant coding, consistent with the idea that multiplexed, high-dimensional representations support efficient spatial computation ^38–40^.

### Comparable hippocampal architecture between species

Spatial coding properties diverged markedly between species (Figs. 3–5), raising the question of whether these differences stem from hippocampal anatomy. The two species differed slightly in hippocampal size and position (Fig. 6a, b), yet cyto- and chemoarchitectonic organization was comparable ^41, 42^. In both species, the CA1 pyramidal layer formed a compact, 50-μm-thick, cell-dense band (Fig. 6c, d), in contrast to the loosely organized primate CA1 ^43^; this organization supports anatomical similarity between rodent and tree shrew hippocampi ^44^. Comparison of our tree shrew single-cell transcriptomic dataset with published rat data ^45^ further revealed similar neuron types across hippocampal regions, with high cross-species concordance (Fig. 6e–g; tree shrew data deposited in NGDC GSA, CRA043801, companion study submitted). Together, these findings are consistent with the view that divergent spatial codes do not require divergent hippocampal architectures, and that representational format can be modified independently of cytoarchitecture and transcriptomic identity.

**Fig. 6.**
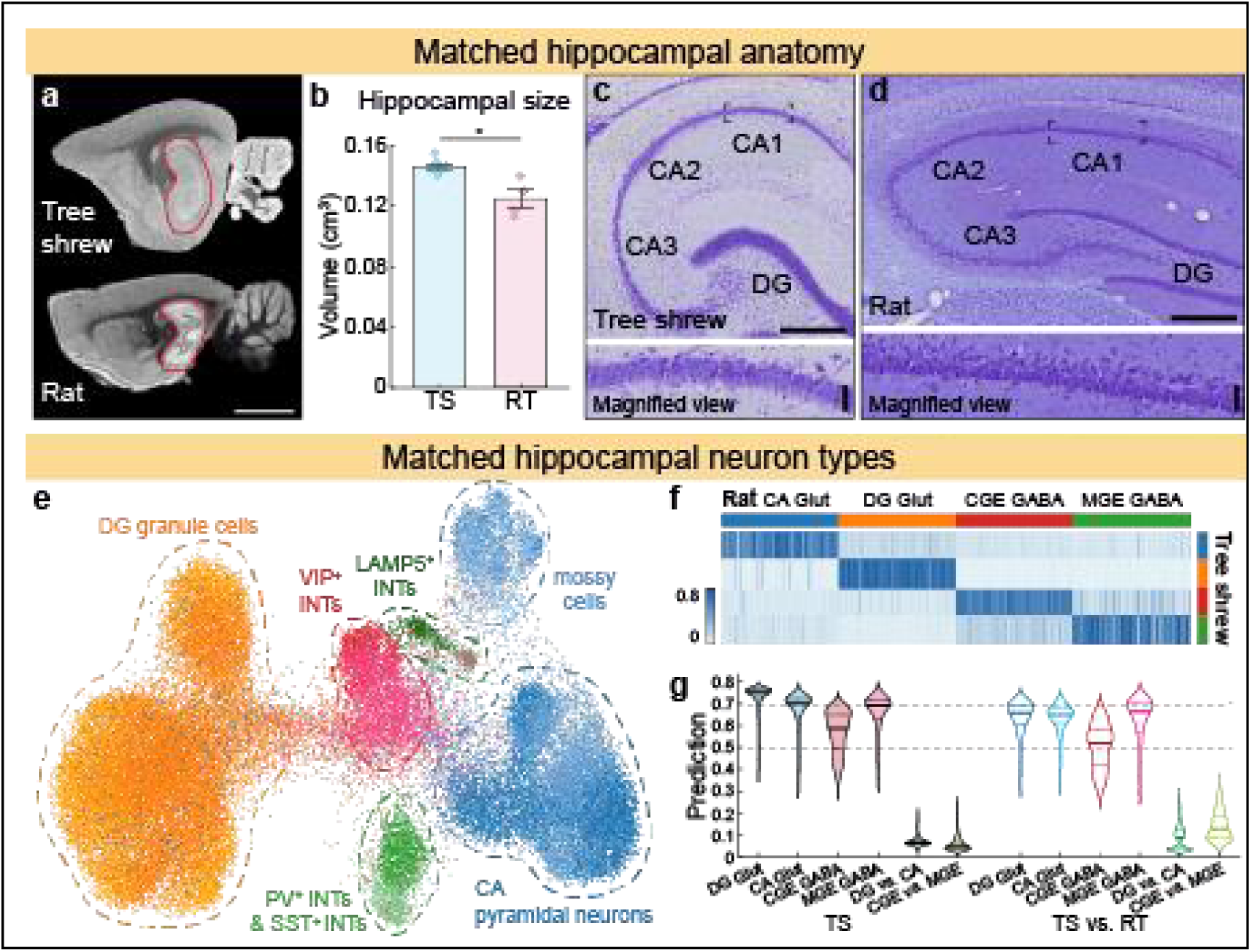
Comparable hippocampal architecture between tree shrews and rats. **a**, Representative MRI sagittal sections. Hippocampus outlined in red. Scale bar: 5 mm. **b**, Hippocampal volume (mean ± SEM). **c**–**d**, Nissl-stained dorsal hippocampus. Top: low magnification; bottom: high magnification of boxed region. DG, dentate gyrus. Scale bars: top, 500 μm; bottom, 50 μm. **e**, Integrated UMAP (Uniform Manifold Approximation and Projection) of hippocampal neuron types. CA, *Cornu Ammonis*; INT, interneuron; LAMP5, lysosome-associated membrane protein 5; PV, parvalbumin; SST, somatostatin; VIP, vasoactive intestinal peptide. **f**, Cross-species cell-type prediction heatmap. Color code as in E; 500 neurons per category. Glut, glutamatergic (excitatory) neurons; GABA, GABAergic (inhibitory) neurons; CGE, caudal ganglionic eminence; MGE, medial ganglionic eminence. **g**, Within-and cross-species prediction scores (median (IQR)). Hippocampal cytoarchitecture and single-cell transcriptomes are highly conserved. \**P* < 0.05.

### Preserved spatial coding across lighting conditions

We tested whether distinct navigational behaviors (Fig. 2) could explain the observed coding differences by recording three tree shrews under normal and dim light. Dim lighting substantially degraded visual input, given the poor scotopic vision of tree shrews ^27^. Consequently, animals shifted from straight transversal movements under normal light to meandering, rat-like trajectories under dim light (Fig. 7a), with increased AV and path tortuosity (Fig. 7b, c; Supplementary Table 1), whereas running speed and spatial coverage remained unchanged (Extended Data Fig. 7a–d).

**Fig. 7.**
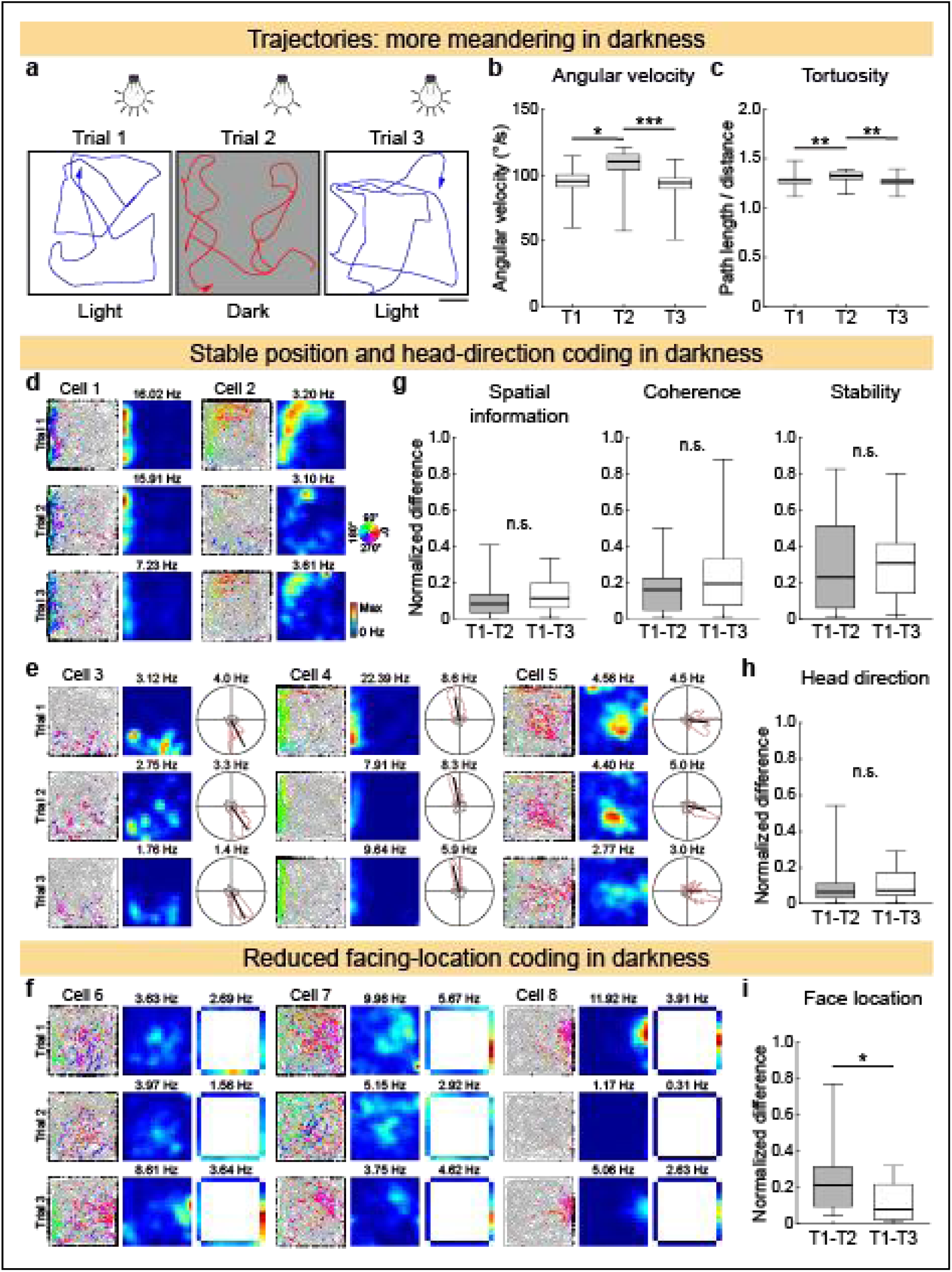
Dim light alters tree shrew navigation but spares spatial coding. **a**, Task design and representative 20-s trajectories. Scale bar: 25 cm. **b**–**c**, Lighting-altered behavioral metrics (median (IQR)). **d**–**e**, Stable position and HD tuning across lighting conditions. **f**, Diminished FL tuning under dim light. Cells 1/3, 2/7, and 5/8 are simultaneously recorded pairs. **g**–**i**, Normalized tuning change indices (median (IQR)). Dim lighting suppresses visual inputs and induces rodent-like meandering, yet the overall spatial coding architecture is preserved. FL neurons are selectively light-sensitive. n.s., not significant; \**P* < 0.05, \*\**P* < 0.01, \*\*\**P* < 0.001.

If weaker position selectivity and stronger multiplexed coding were driven by vision-guided straight-path locomotion, we would expect enhanced position tuning and diminished HD and FL tuning under dim light. Contrary to this prediction, most spatial neurons retained stable firing patterns across lighting conditions (Fig. 7d, e), with consistent overall spatial tuning (Extended Data Fig. 7e–l), in agreement with analogous rodent experiments ^46, 47^. These results argue against the hypothesis that coding differences are dictated by locomotor behavior. A small subset of neurons exhibited dramatic reductions in spatial firing between normal-light and dim trials (Fig. 7f), whereas simultaneously recorded cells remained largely unaltered (Fig. 7d, e, cells 2 and 5). These light-sensitive neurons were predominantly FL-selective. Normalized tuning changes differed significantly in FL neurons between lighting conditions (T1–T2) versus repeated trials under identical conditions (T1–T3), whereas position and HD tuning remained intact (Fig. 7g–i). These observations are consistent with SV-like coding in tree shrews, analogous to primates, in which FL serves as a close proxy for light-dependent SV coding ^5, 6, 9, 48^.

Collectively, dim lighting suppressed visual input and induced meandering, rat-like exploratory behavior. Yet this manipulation did not shift hippocampal representations toward a rodent profile: the overall architecture of spatial coding was preserved despite the degraded sensory conditions, a pattern-completion–like process ^49^. These results suggest that inter-species coding differences reflect evolutionary divergence in computational strategy rather than direct behavioral adaptation to sensory inputs.

### Global remapping with preserved network structure in tree shrew CA1

Rodent place cells orthogonalize activity across contexts via global remapping ^15^, as do monkeys ^50^. We examined whether tree shrew CA1 exhibits analogous remapping by recording in identical arenas across spatially distinct rooms. Nine animals were tested in the two-room task, with trials 1 and 3 in one room and trial 2 in another. Most neurons remained active in both rooms (Fig. 8a). Position-modulated neurons showed stable firing between repeated same-room trials but shifted or ceased firing across environments, consistent with global remapping ^51^ (Fig. 8b). Non-spatial neurons fired randomly regardless of environmental layout (Fig. 8b, tree shrew cell 7).

**Fig. 8.**
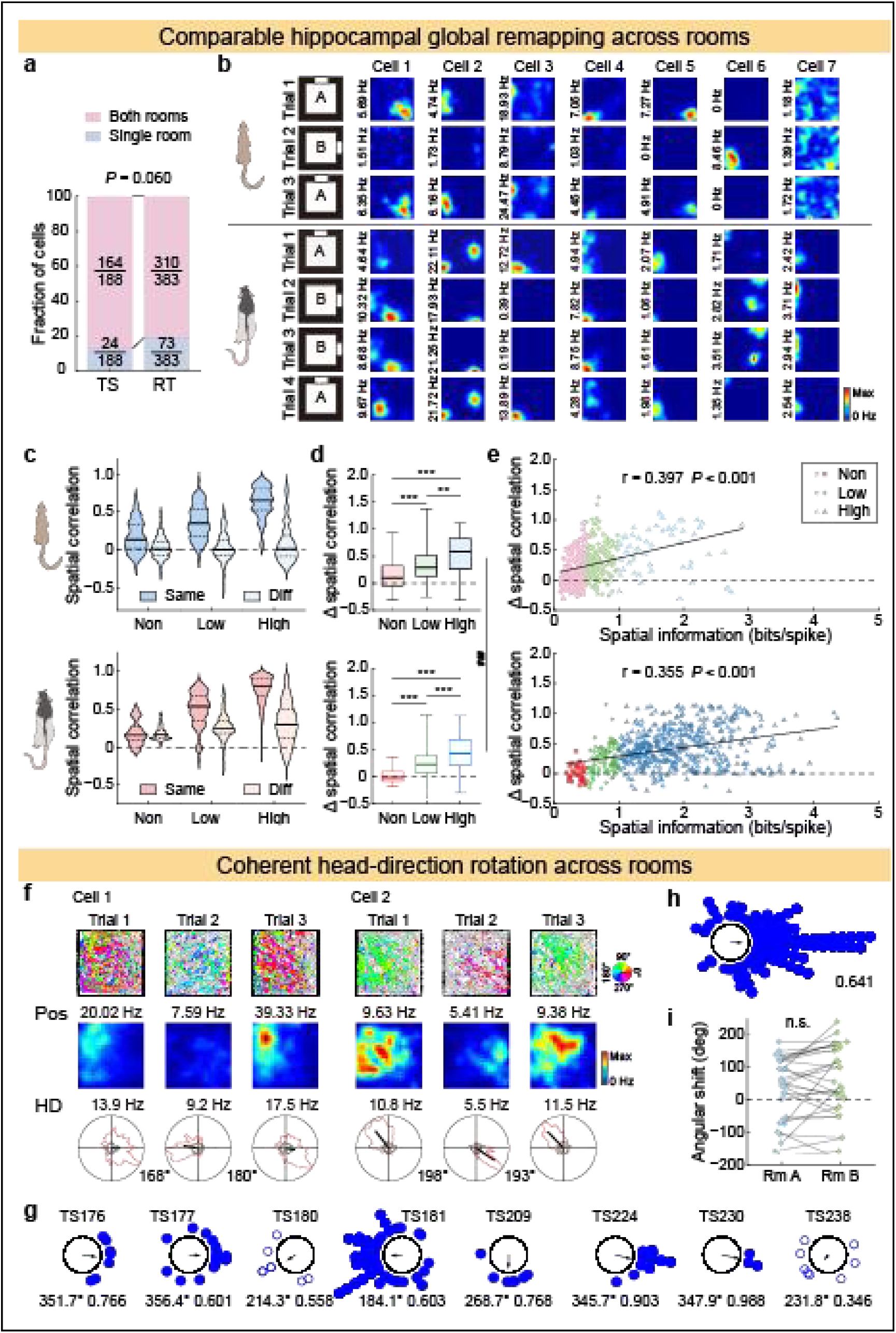
Global remapping with preserved network structure in tree shrew CA1. **a,** Fraction of CA1 neurons active in one or both rooms. **b**, Representative rate maps in two distinct rooms (schematic left). **c**, Spatial correlation (SC) distributions by selectivity group (median (IQR)). **d**, Room-switch ΔSC (outliers hidden). **e**, SIc vs. ΔSC. Solid line, significant regression. **f**, Representative neurons showing coherent ∼180° HD rotations across rooms. **g**, Direction shift distributions across rooms for HD neurons in each tree shrew. Dots (circles when non-significant), individual neurons; central arrows/numbers, mean angular shifts and MVLs. **h**, Normalized direction shifts (raw minus per-animal median) all HD-tuned neurons. **i**, Pairwise HD angular differences for simultaneously recorded HD neuron pairs across rooms. Each point represents the mean across all HD pairs within one experiment. Tree shrew CA1 orthogonalizes activity across environments and preserves network relationships. Cross-group: \**P* < 0.05, \*\*\**P* < 0.001; cross-species: ^###^*P* < 0.001.

Because recordings did not span the entire CA1 transverse axis, we sorted active excitatory neurons by SIc into non-positional (≤ 0.5 bits/spike) ^52^, low-positional (0.5–1.0 bits/spike), and high-positional (≥ 1.0 bits/spike; cutoff chosen for this study) groups. We compared spatial map similarity across trials via spatial correlation (SC). All tree shrews exhibited global remapping (Extended Data Fig. 8a): SC values approached zero across rooms for all groups, indicating near-complete map reorganization. In contrast, same-room SC values increased gradually from non-to high-positional neurons (Fig. 8c, top), producing graded differences between cross-room and same-room similarity (Fig. 8d, top) that correlated significantly with individual position selectivity (Fig. 8e, top). Rats showed a similar trend but with significantly higher spatial tuning and within-room similarity (species main effect, *P* < 0.001; species × group, *P* = 0.332; Fig. 8c–re, bottom). Comparable gradients were obtained using rate difference and population vector correlation analyses (Extended Data Fig. 8b–g), confirming graded environmental discrimination across groups.

Network structure is thought to be preserved across remapping by entorhinal inputs, despite apparently random remapped firing locations ^53^. We therefore examined whether HD shifts were coherent across environments in the tree shrew hippocampus. Preferred HDs shifted coherently across rooms in individual animals (Fig. 8f; Supplementary Table 1): in six of eight tree shrews with HD cells, HD neurons rotated in ∼90° steps, suggesting geometric boundary influence (Fig. 8g). Normalized HD shifts (raw minus mean per animal) showed low dispersion across neurons (Fig. 8h), and angular differences between simultaneously recorded HD neuron pairs were retained across rooms (Fig. 8i). These observations indicate preserved network relationships in the tree shrew hippocampus across spatial environments, analogous to rodent medial entorhinal cortex (MEC) ^54, 55^.

Together, position-selective neurons in tree shrew CA1 orthogonalize activity across environments, as in rodents and primates, while network organization is preserved, likely constrained by upstream entorhinal inputs. These findings indicate conserved hippocampal responses to environmental change across mammalian species.

## DISCUSSION

The mammalian hippocampus supports diverse representational formats across species, yet the transition from position-dominant rodent coding to multiplexed primate coding has remained unexplored. Using identical methods across species, we establish tree shrews as a phenotypic intermediate in hippocampal spatial coding: CA1 neurons exhibited moderate position selectivity alongside multiplexed coding approaching primate characteristics, together with conserved proximo-distal organization and preserved ancestral network dynamics including pattern completion and global remapping. Notably, these hybrid properties were neither a consequence of distinct hippocampal architecture nor directly driven by navigational behavior. Multiplexed populations achieved rat-comparable decoding accuracy with greater efficiency, suggesting computational advantages of high-dimensional coding. These findings support a quantitative transition from position-dominant to multiplexed spatial representations consistent with quantitative repurposing of entorhinal-hippocampal circuitry, rather than qualitative architectural innovation (Extended Data Fig. 9).

This intermediate phenotype is first evident in the composition of spatial representations. Tree shrew CA1 exhibits a coding profile between rodent position-dominant and primate view/orientation-dominant coding ^5–8^, with position-selective neurons comprising 87–95% of spatial cells in rats, 44–69% in tree shrews, and 14–32% in monkeys ^5, 6, 9, 17, 29, 32^, whereas FL/SV-coding neurons comprise 10–19% in tree shrews versus 18–22% in macaques ^5, 7^. We found that dim lighting selectively impairs FL neurons, substantiating their visual dependence and mirroring primate SV cell properties ^48^. Moreover, pronounced HD tuning may similarly reflect visual reliance, which strengthens HD modulation in virtual environments ^56^, and in visually oriented rodents ^57^. Tree shrews are diurnal, with laterally positioned eyes ^58^ and rodent-like coordinated eye-head rotation ^59^, yet their visual capabilities exceed nocturnal rodent standards while lacking primate foveal precision ^27, 60–64^. This intermediate visual ecology aligns with their hippocampal coding: stronger HD and FL/SV representation than rodents, but substantially less visual dominance than primates. Primates use high-resolution vision for look-ahead navigation, sampling distant landmarks without direct visitation; rodents, constrained by poor acuity, must physically locomote to sample spatial relationships. Tree shrews occupy a functional middle ground, supporting more extensive visual spatial sampling than rodents without primate-style foveal strategies.

This hybrid phenotype challenges three prevailing place cell models. The boundary-vector model predicts that sharper directional and egocentric inputs should yield more confined place fields ^65, 66^, yet tree shrew fields are more dispersed. The retinal-resolution model predicts that higher acuity should enable sharper single-cell position tuning ^67^, yet tree shrews show weaker position selectivity. The view-field model predicts that the wide visual fields shared by rodents and tree shrews should favor rodent-like place fields ^68^, yet tree shrews exhibit stronger FL modulation. These discrepancies, together with comparable decoding accuracy from multiplexed populations using fewer neurons, indicate that position selectivity depends less on sensory parameters than on the computational strategy for integrating visual and self-motion cues ^13^. Tree shrews appear to sacrifice positional precision for flexibility, prioritizing multiplexed representations of HD and viewing angle over rigid boundary anchoring, an arboreal specialization consistent with theoretical frameworks for integrated neural representations ^38–40^.

Despite divergent spatial coding schemes, fundamental network properties remain conserved between rats and tree shrews, with several anatomical and computational features similarly preserved in primates. Our light manipulation shows that tree shrew hippocampal coding remains largely stable when visual inputs are suppressed, paralleling rodent place cell persistence in darkness, albeit with reduced accuracy ^46, 47^. This stability indicates a preserved network mechanism that maintains established cognitive maps from degraded sensory inputs, suggesting that attractor-like dynamics are a fundamental property of mammalian hippocampal networks enabling continuous localization when sensory cues become unreliable ^49^.

Tree shrew CA1 likewise exhibits global remapping, orthogonalizing activity patterns across distinct environments as in rodents and primates ^15, 50^, indicating preserved computation despite quantitative differences in tuning strength. The coherent, geometrically constrained HD rotations observed in tree shrews mirror the rigid spatial organization of entorhinal grid, border, and HD cells in rats ^54, 55, 69^, suggesting that upstream circuits constrain hippocampal remapping through conserved spatial reference frames ^53^. In both species, position selectivity correlated with sensitivity to spatial change, perhaps enhanced by realignment of MEC spatial cell activity^54, 70^.

The proximo-distal gradient of spatial variables in both tree shrews and rats further indicates that functional segregation along the CA1 transverse axis represents an ancestral mammalian trait, potentially reflecting conserved differences in entorhinal inputs ^71–73^. Preferential proximal coding of allocentric variables, including position, HD, and FL, likely reflects strong MEC inputs from position- and HD-selective neurons ^74^. Conversely, enhanced distal coding of EB in first-order models (Extended Data Fig. 6a) may arise from stronger lateral entorhinal inputs conveying egocentric information ^75^. Given the conserved medial-lateral entorhinal dichotomy from rodents to primates ^71, 76^, the proximo-distal gradient likely represents a deeply preserved organizational principle, with segregated entorhinal inputs maintaining consistent computational roles across species, despite circuit expansion in larger brains ^76^.

Together, these findings indicate that the entorhinal-hippocampal circuit operates as a conserved computational scaffold whose representational format can be tuned by upstream modifications without altering core network dynamics. Species-specific demands shape the statistical structure of cortical inputs to the entorhinal cortex ^73, 76^, determining whether coding is position-dominant or multiplexed; the hippocampus repurposes ancestral dynamics rather than evolving de novo representations. This principle is supported by anatomical homology between tree shrew and rodent hippocampi ^41, 42, 44^ and conserved entorhinal-hippocampal projections across species ^73, 76–78^. Notably, anatomical comparability in this study is limited to cytoarchitecture and cell-type composition; dendritic morphology and neuromodulation remain unexamined and could likewise contribute to coding divergence. Quantitative differences in spatial representation may reflect progressive reorganization of MEC layer II, from alternating pyramidal clusters and stellate neurons in rodents, through an intermediate salt-and-pepper distribution in tree shrews, and stratified tiers in monkeys ^78–80^, which may alter the temporal dynamics of afferent integration. Such microcircuit modifications within a conserved macroscopic architecture may provide a mechanistic basis for the evolutionary transition from rodent-like to primate-like spatial coding, although direct causal evidence remains to be established.

The hybrid representational phenotype and conserved network dynamics described above allow us to adjudicate among the three explanations for rodent-primate divergence. First, applying identical tasks, environments, and analytical tools, we ruled out behavioral demands as the primary source of hybrid coding: both species were tested in identically sized arenas with matched cue configurations using the same free-navigation task, and were trained to sample space uniformly. Critically, position and HD coding remained largely unaltered across lighting conditions in tree shrews, indicating that representational format does not simply track sensory availability or navigational strategy. Second, conserved cytoarchitecture and overlapping neuron types between rat and tree shrew hippocampi argue against qualitative architectural innovation. The further preservation of fundamental network dynamics, including pattern completion and global remapping, indicates that the entorhinal-hippocampal scaffold is not radically reorganized in this lineage. Third, weaker position tuning and enhanced multiplexed coding within this conserved architecture support flexible optimization of coding strategy, specifically, quantitative repurposing of ancestral dynamics. Together, these observations indicate that the shift from rodent-like to primate-like coding can occur through incremental representational change without wholesale circuit redesign.

These findings lay the groundwork for investigating the evolutionary and ecological dimensions of spatial cognition. However, phylogenetic position and sensory ecology are inherently confounded in a single intermediate species: our data establish the phenotype’s intermediate position but cannot determine whether it reflects shared ancestry with primates or convergent ecological adaptation. Dissociating these factors will require additional species at off-diagonal positions, such as squirrels, diurnal, arboreal rodents phylogenetically distant from primates ^27^. Characterizing entorhinal cortex function in tree shrews is another priority, clarifying whether hippocampal multiplexing arises from transformed entorhinal inputs or emerges through local computation. Causal manipulations, such as targeted inactivation of visual cortex or entorhinal subregions, could test whether visual dependence reflects obligatory processing or flexible circuit specialization. Finally, tree shrews’ closer phylogenetic relationship to primates than rodents may refine preclinical translation of spatial memory disorders.

## RESOURCE AVAILABILITY

### Lead contact

Further information and requests for resources and reagents should be directed to and will be fulfilled by the Lead Contact, Li Lu.

### Materials availability

This study did not generate new unique materials.

### Data and code availability

Tree shrew single-cell transcriptomic data are deposited in NGDC GSA (CRA043801; to be released upon publication of the companion study). Custom analysis code is available as Supplementary Code. Rat single-cell transcriptomic data as well as standard algorithms and publicly available software tools are listed in the Key Resources Table. Other data supporting the findings of this study are provided in the Supplementary Data files. Additional datasets are available from the corresponding author upon reasonable request.

## SUPPLEMENTARY INFORMATION

Extended Data Figs. 1 – 9, Supplementary Tables 1 and 2, and Supplementary Video 1 are available online.

## ACKNOWLEDGMENTS

This research was supported by the Brain Science and Brain-like Intelligence Technology - National Science and Technology Major Project (2022ZD0205000 to L.L., B.L.S. and C.R.L.; 2021ZD0200900 to Y.G.Y.), and “Light of West China” Program of the Chinese Academy of Sciences (xbzg-zdsys-202404 to L.L. and C.R.L.; xbzg-zdsys-202302 to Y.G.Y), Yunnan Revitalization Talent Support Program (Yunling Scholar Project to L.L.), Yunnan Province (202305AH340006 to Y.G.Y.) and National Natural Science Foundation of China (W2512020 and 32427802 to C.R.L.).

The authors express their gratitude to Cheng-Ji Li, Rong Zhang, Ning Xu, Rui Bi, as well as the staff members of the National Research Facility for Phenotypic & Genetic Analysis of Model Animals (Primate Facility) (https://cstr.cn/31137.02.NPRC), for providing technical support and assistance in data collection and analysis. They also thank Xin-Jian Li for his valuable suggestions on the manuscript.

## AUTHOR CONTRIBUTIONS

L.L. conceived and designed the study. X.F.G. established the wireless recording methodology in tree shrews and, together with C.D., Y.Q.H., and Y.F.Y., performed the tree shrew recording experiments. C.D., X.F.G., Y.L.D., S.Y.H., and Q.C.X. performed the neural data analyses. J.W.Z., W.C.L. and B.L.S. decoded the neural data. K.J. and C.R.L. collected and analyzed the MRI data. W.B.K. and Y.G.Y. analyzed the single-cell transcriptome data. C.D., X.F.G., J.L.L., and L.L. prepared the figures. X.C., and L.L. wrote the manuscript. All authors reviewed and edited the manuscript.

## Methods

### Ethical compliance

All procedures were designed to minimize animal suffering and reduce the number of subjects required. Experiments were approved by the Institutional Animal Care and Use Committee (IACUC) of the Kunming Institute of Zoology (Approval Nos.: KIZ-IACUC-TE-2021-10-001 for tree shrews; KIZ-IACUC-RE-2021-06-019 for rats) and conducted in accordance with national regulations and institutional guidelines for laboratory animal care and use.

### Animal models

Twenty-one adult male tree shrews (*Tupaia belangeri chinensis*; 17–42 months old; 120–180 g) and twenty-four adult male Long-Evans rats (19–34 weeks old; 500–600 g) were used. Fourteen tree shrews (25.1 ± 1.9 months old) and twenty rats (26.9 ± 1.1 weeks old) underwent recording experiments. Animals were comparably age-matched, given that tree shrews live 3 to 4 times as long as rats (6–8 years vs. ∼2 years) ^25^. The rat recording data were previously published ^19^; however, the present study employs distinct analytical methods.

Drive-implanted animals were housed individually (tree shrew: 40 × 38 × 35 cm; rat: 46 × 30 × 36 cm, W × D × H) under standard laboratory conditions (temperature: 20–23°C; humidity: 40– 60%; 12-h light/dark cycle). Light schedules were species-specific: tree shrews were diurnal (lights off 19:00–07:00); rats were nocturnal (lights off 09:00–21:00). Behavioral testing occurred during each species’ active period (tree shrews: light phase; rats: dark phase). During habituation, handling and post-surgery recovery, animals had *ad libitum* access to food and water. During recording experiments, they were mildly food-restricted to 85–90% of their free-feeding body weight.

### Electrode preparation

#### Tree shrews

Custom-built drive assemblies consisted of two independently movable bundles, each containing four tetrodes.

#### Rats

Multi-tetrode “hyperdrives” comprising 18 independently movable tetrodes ^81^ were used. Tetrodes were constructed from 17-μm-diameter polyimide-coated platinum-iridium wire (90:10 Pt-Ir; California Fine Wire, USA). Prior to implantation, tetrodes were electroplated to achieve impedances of 150–250 kΩ measured at 1 kHz (Biomega; Bio-Signal Technologies, China). All drive assemblies were gas-sterilized with ethylene oxide (Sanqiang Medical, China) within 48 h prior to surgery.

### Surgery

Anesthesia was induced with isoflurane (0.5–3% v/v in oxygen, 0.8–1.0 L/min; RWD Life Science, China) and maintained at concentrations adjusted according to physiological monitoring (respiratory rate, withdrawal reflexes). Body temperature was maintained at 38°C with a feedback-controlled heating pad (RWD Life Science, China). Preoperative medications included meloxicam (rat: 1 mg/kg; tree shrew: 5 mg/kg), enrofloxacin (5 mg/kg), and atropine (rat: 0.1 mg/kg; tree shrew: 0.5 mg/kg). The scalp was shaved, disinfected, and locally anesthetized with 2% lidocaine. Ophthalmic ointment was applied to prevent corneal desiccation. Animals were secured in a stereotaxic frame (RWD Life Science, China) with the skull leveled (bregma and lambda aligned horizontally).

#### Tree shrews

Bilateral craniotomies were performed at AP 4.0–4.7 mm and ML ±5.2–5.6 mm relative to bregma. The drive was gradually lowered to insert tetrode tips ∼3 mm into the cortex overlaying both dorsal hippocampi.

#### Rats

Tetrodes were inserted ∼1 mm into the cortex above the right hippocampus, at AP 3.2–5.6 mm and ML 1.0–4.5 mm relative to bregma.

Two stainless-steel screws implanted above the cerebellum served as ground. The drive was secured to the skull using additional stainless-steel screws and dental acrylic cement. Postoperative analgesia (meloxicam) and antibiotics (amoxicillin) were administered daily until full recovery. Recording experiments commenced after ≥ 1 week of recuperation.

### Electrophysiological recordings

During the 3–4 weeks following surgery, tetrodes were advanced ventrally in daily increments of ≤ 40 μm until high-amplitude complex-spike activity indicated positioning within the hippocampal CA1 pyramidal cell layer ^82^. Electrophysiological recording procedures followed previously described methods ^19^. Recordings commenced only after signals remained stable at the target location for ≥ 24 h. Repeated sampling from the same tetrode was accepted only if subsequent recording sites were separated by ≥ 40 µm, minimizing overlap between cell populations.

#### Tree shrews

Broadband neural signals (0.3–7 500 Hz) were recorded using a 32-channel wireless neural logger (Hermes; Bio-Signal Technologies, China). Signals were amplified (×192), digitized at 30 kHz by a 16-bit analog-to-digital converter (resolution: 0.15 μV), and stored on a 32 GB microSD card. The system was powered by a lightweight 3.7 V Li-Po battery. The logger was wirelessly synchronized with the computer clock every 5 s via an infrared transceiver. Differential recordings were referenced to two cerebellar skull screws.

#### Rats

Unit activity and local field potentials (LFPs) were recorded using a 64-channel wired data acquisition system (Zeus; Bio-Signal Technologies, China). Unit activity was amplified using the same RHD2000 amplifier chip (Intan Technologies, USA) as in tree shrews, band-pass filtered (300–7 500 Hz, 3^rd^-order Bessel filter), and digitized at 30 kHz (16-bit resolution, 0.15 μV). Spike waveforms exceeding a −50 μV threshold were time-stamped and recorded for 1 ms. LFPs were acquired simultaneously from one channel per tetrode (0.3–300 Hz band, 1 kHz sampling rate), referenced to the two cerebellar skull screws.

### Test environments

Recording arenas (100 × 100 × 50 cm; W × D × H) consisted of metal frames fitted with four black plastic walls. A single white cue card (30 × 50 cm; W × H) was centered on one wall to maximize salience. Distal visual cues were eliminated by floor-to-ceiling curtains (120 × 120 cm). A resting box was positioned outside the curtains.

The resting box was positioned outside the curtains between the arena and the experimenter.

Lighting conditions were species-specific: rats were tested under dim illumination (< 10 lux); tree shrews under standard laboratory lighting (∼50 lux), except during the dark condition of the light-dark task (< 10 lux). Experimental setups were standardized across species; notably, many recordings for both species were conducted in the same testing room. For the two-room task, two distinct testing rooms of equivalent size but different spatial configurations were employed. Recording boxes were visually identical in both rooms but differed in cue card position.

### Behavioral paradigms

Tree shrews were trained to forage for randomly scattered cake crumbs ^83^ until they consistently explored > 90% of the arena within the trial duration. Subjects that failed to complete more than one trial were excluded from subsequent testing. Experiments consisted of either single 8–15 min trials or three-trial sessions (A-B-A design) with 8-min trials separated by 5-min inter-trial rests (ITRs) in the resting box. A-B-A tasks included two-room remapping task (A: room A; B: room B) and light-dark manipulation task (A: standard illumination; B: reduced illumination). The arena floor was cleaned with 75% ethanol after each trial.

Two-room task: trials 1 and 3 were conducted in Room A and trial 2 in Room B. Animals were TS176, TS177, TS180, TS181, TS195, TS209, TS224, TS230 and TS238; all exhibited clear global remapping. Light-dark task: trials 1 and 3 were conducted under standard illumination (∼50 lux), whereas trial 2 was under dim illumination (<10 lux), with unchanged arena geometry and room layout. Animals were TS230, TS238 and TS240.

Rats were trained to forage for randomly scattered cookie crumbs for 2–3 weeks until they consistently explored > 90% of the arena within 10 min, and for multiple trials. Detailed methods for rat behavioral testing have been previously described ^19^; briefly, sessions followed an A-B-B-A design (four 10-min trials per session) with 5-min ITRs. Conditions A and B denoted distinct manipulations of room layout (spatial) or wall color (non-spatial). The floor was cleaned with 75% ethanol during ITRs. Six rats that failed to exhibit global remapping in the two-room task were excluded from subsequent global remapping analyses.

### Histology and recording site localization

Following recordings, animals were deeply anesthetized and transcardially perfused with 0.9% saline followed by 4% formalin, leaving tetrodes in place. Brains were extracted, post-fixed for 24 h, cryoprotected in 30% sucrose, and sectioned at 40 μm (coronal for rats, sagittal for tree shrews) with a cryostat (CT520; Dakewe, China). Brain sections were Nissl-stained with cresyl violet (CAS 10510-54-0, Cat. no. C5042-10g; Sigma-Aldrich, USA) and captured in arrays of FOVs at a resolution of 0.69 μm per pixel under a bright-field microscope (BX61, 10×/0.4 NA objective; Olympus, Japan; RRID: SCR_020343), then reconstructed and corrected for shading artifacts using OlyVIA software v3.2 (Olympus, Japan; RRID: SCR_016167).

Hippocampal subregions were delineated using established cytoarchitectonic criteria ^16, 41, 73^. Only data from tetrodes confirmed to be located within the CA1 pyramidal cell layer were included. The proximo-distal position of each site was quantified by measuring the distance from the CA1/CA2 boundary to the tetrode tip with ImageJ (v1.52a; NIH, USA; RRID: SCR_003070), normalized to the total transverse length of CA1 (CA2 border = 0; subiculum border = 1). Sites were classified as proximal (pCA1; 0–0.333), middle (mCA1; 0.333–0.667), or distal (dCA1; 0.667–1.0) relative to the CA2 border (0) and subiculum (1).

### Magnetic resonance imaging (MRI)

Seven male tree shrews (∼3 years old) and four male rats (∼5 months old) underwent *ex vivo* MRI. Animals were euthanized with CO₂ and transcardially perfused with 0.9% saline followed by 4% paraformaldehyde (PFA) in 0.1 M phosphate buffer. Brains with the skull intact were immersion-fixed in 4% PFA containing 0.3% gadolinium contrast agent for 2–4 weeks, then rinsed and stored in phosphate-buffered saline (PBS) for one week prior to imaging. High-resolution MRI was performed on a 9.4 T Bruker BioSpec 94/30 scanner (Bruker, USA) using a fast low-angle shot (FLASH) sequence.

#### Tree shrews

(T1-weighted): time of repetition (TR) = 20 ms, time of echo (TE) = 5.8 ms, matrix = 440 × 660 × 240, field of view (FOV) = 22 × 33 × 18 mm³, flip angle = 20°, voxel size = 50 × 50 × 75 µm³, and scan time = 29 min 41 s.

#### Rats

(T2-weighted): TR = 30 ms, TE = 20 ms, matrix = 200 × 280 × 180, FOV = 20 × 28 × 18 mm³, flip angle = 25°, voxel size = 100 × 100 × 100 µm³, and scan time = 14 min 17 s.

Whole-brain and bilateral hippocampal volumes were quantified by manual segmentation using ITK-SNAP (v4.4.0; http://www.itksnap.org/pmwiki/pmwiki.php) and FSL (v6.0.4; https://fsl.fmrib.ox.ac.uk/fsl/docs/index.html) software packages ^84, 85^.

### Cross-species single-cell alignment and cell-type annotation

Publicly available rat (NCBI GEO: GSE306235) and tree shrew (NGDC/GSA: CRA043801; to be released upon publication of the companion study) scRNA-seq datasets were processed using Scanpy ^86^ (https://github.com/scverse). Initial cell-type identities were established by mapping both datasets against the Allen Brain Atlas via the MapMyCell portal (https://knowledge.brain-map.org/mapmycells/process). Cross-species data integration and cell-type projection were then performed using CAME (cell-type assignment and module extraction; https://github.com/liuxiaoyu-git/CAME) ^87^ to generate a unified comparative landscape. Uniform manifold approximation and projection (UMAP) was applied to the joint embeddings for visualization of cross-species alignment and topological cell distributions. Cell-type assignment probabilities generated by CAME were projected onto UMAP coordinates to evaluate mapping confidence.

### Spike sorting and cell classification

Tree shrew raw neural signals were high-pass filtered at 300 Hz (Bessel filter), and spikes from each tetrode were detected by threshold crossing (−50 µV) to match the sampling of rat recordings. Detected waveforms (1 ms duration) were time-stamped and extracted using MATLAB (MathWorks, USA; RRID: SCR_001622). For both species, single units were isolated offline using MClust (v4.4; https://github.com/adredish/MClust-Spike-Sorting-Toolbox) by examining 2D projections of spike features (peak amplitude, energy, peak-to-valley amplitude) across all four tetrode channels. Cluster quality was validated using waveform shape and autocorrelation based on spike timing. Spike sorting was conducted on concatenated data from the entire recording experiment; spike trains were subsequently segmented by trial.

Putative excitatory pyramidal neurons were identified using three established criteria: (1) spike width (peak-to-trough) ≥ 200 µs; (2) mean firing rate ≤ 2 Hz; and (3) presence of occasional complex-spike bursts ^51^. Units that failed any criterion or that were recorded repeatedly in the same task (defined by similar spatial firing properties and/or inter-spike interval distributions) were excluded.

### LFP power analysis

Tree shrew LFPs were obtained by low-pass filtering raw neural data at 500 Hz (4th-order Chebyshev Type II) and down-sampling to 1 kHz to match the sampling rate of rat recordings. For both species, a 50 Hz notch filter was applied to eliminate mains frequency interference. Time-frequency decomposition for visualization used the continuous wavelet transform. Absolute power spectra (0–100 Hz) were estimated from consecutive 5-s windows using the multitaper method (Chronux v2.12; http://chronux.org; time-bandwidth product = 3, 5 tapers) and the Buzsáki lab toolbox (https://github.com/buzsakilab/buzcode). Aperiodic (1/f-like) components were parameterized and removed using the FOOOF algorithm ^31^ (https://github.com/fooof-tools/fooof) to isolate oscillatory power and control for broadband noise differences across recording systems. Experiments with poor LFP quality were excluded.

### Theta phase locking

Theta-phase locking was assessed by quantifying the coupling between CA1 unit activity and hippocampal theta oscillations (7–10 Hz). Instantaneous theta phases were derived using the Hilbert transform using the MATLAB Signal Processing Toolbox. Each spike was assigned the nearest theta phase value at its corresponding time point. Phase values were then binned into 10° intervals (n = 36 bins) spanning 0–360°.

Phase locking strength was characterized by the mean vector length (MVL) using the Circular Statistics Toolbox ^88^ (v1.21; https://github.com/circstat/circstat-matlab):

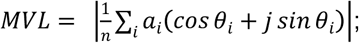

where *j* represents the imaginary unit, *n* is the number of phase bins (36), and *θ_i_* and *a_i_* denote the theta phase and normalized response (spike count) of the *i*-th bin, respectively.

### Spatial variable extraction

Raw position data were tracked at 50 Hz using an overhead camera and head-mounted red and green light-emitting diodes (LEDs) for both species. The *x*- and *y*-coordinates of red and green LEDs were smoothed with a 0.5-s sliding window. The following variables were derived:

Tracking used an overhead Cyclops camera system (Bio-Signal Technologies, China).

#### Position (Pos)

Average of red and green LED coordinates in the horizontal plane.

#### Head direction (HD)

Orientation of the naso-occipital axis in the horizontal plane, computed from the vector connecting the two LEDs (forward = head centroid to nose).

#### Facing location (FL)

Allocentric coordinates of the intersection point between the arena boundary and the ray extending from the head position along the HD vector (i.e., the wall location where the animal is facing). This variable serves as a proximal measure of spatial view commonly studied in primates ^5, 6^.

#### Egocentric bearing (EB)

Angle between the HD vector and the vector connecting the head center to the arena center, measured in a head-centered reference frame (0° = facing center)_34._

#### Linear speed (LS)

Translational speed calculated as Euclidean displacement over a ±20-ms sliding window.

#### Angular velocity (AV)

Rate of change in HD (°/s), calculated as the angular difference between consecutive samples divided by the sampling interval (20 ms).

### Behavioral metrics

Head position was tracked online at 50 Hz using head-mounted LEDs during freely behaving recording sessions. Seven metrics quantified navigational behavior across trials.

#### Median speed (LS)

Median instantaneous translational speed (cm/s) within each trial.

#### Median AV

Median absolute head angular velocity (°/s) within each trial.

#### AV/LS ratio

Median AV divided by median speed (°/cm) within each trial.

#### Path tortuosity

Local tortuosity measured as the ratio of actual path length to straight-line distance between trajectory endpoints, calculated within a ±1-s sliding window and then averaged across the trial ^89^. Values range from 1 (straight-line movement) to > 1 (circuitous paths).

#### Thigmotaxis

The ratio of time spent in the arena periphery (border zone, 15 cm from walls) to time spent in the center (70 × 70 cm) during exploration.

#### Spatial coverage

Proportion of 5 × 5 cm spatial bins visited for > 0.5 s, expressed as a percentage of total arena bins.

#### Spatial evenness

Mean global Moran’s *I* quantifying the spatial autocorrelation of occupation time across the arena. The index was calculated as:

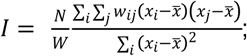

where *N* is the total number of spatial bins (indexed by *i* and *j*), *w_ij_* is the spatial weight between bins *i* and *j* (second-order queen’s contiguity matrix), *W* is the sum of all spatial weights, *x* and *x̃* are the occupation time in each bin and the mean occupation time across all bins, respectively. Values approaching 0 indicate random spatial exploration; values approaching +1 indicate strong clustering (repetitive sampling of specific locations).

### Spatial rate maps and place fields

Data from periods of immobility or tracking artifacts were excluded based on speed thresholds (< 2.5 cm/s for rats, < 5 cm/s for tree shrews; or > 100 cm/s). Only neurons with mean firing rates ≥ 0.1 Hz (total spikes divided by trial duration) were included. Spatial rate maps were constructed by binning spike counts into 5 × 5 cm spatial bins, dividing by dwell time per bin, and smoothing with a 2D Gaussian kernel (σ = 1 bin). Place fields were identified as contiguous regions meeting three criteria: (1) minimum area of 225 cm² (≥ 9 adjacent bins); (2) firing rate in constituent bins ≥ 20% of the cell’s peak rate; and (3) peak firing rate within the field ≥ 1 Hz ^19^. Distinct, non-overlapping fields were enumerated separately for each cell.

### Spatial modulation

spatial information rate (bits/second): 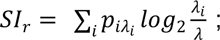;

spatial information content (bits/spike): 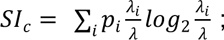;

where *λ_i_* is the mean firing rate in the *i*-th spatial bin, *λ* is the overall mean firing rate of the cell, and *p_i_* is the occupancy probability for that bin (time in bin *i* divided by total trial time). Spatial coherence: Pearson correlation between firing rate in each unsmoothed spatial bin and the mean firing rate of its eight immediately adjacent bins (first-order spatial autocorrelation). Spatial stability: Pearson correlation between smoothed rate maps generated from the first and second halves of each trial.

### Tuning curves

Tuning curves for circular variables (HD, FL, and EB) were generated by assigning spike counts to 3° angular bins, divided by dwell time, and smoothed with a 1D Gaussian kernel (σ = 1 bin). Tuning strength and preferred orientation were characterized by the MVL and the circular mean angle of the distribution, respectively.

### Permutation testing

Spatial metrics: significance of spatial modulation was assessed via permutation testing ^16^. For each cell, 1,000 surrogate datasets were generated by circularly time-shifting the spike train relative to the behavioral trajectory by a random interval (20 s to trial duration minus 20 s), with trial boundaries wrapped to maintain data continuity. This procedure was applied to all CA1 cells recorded from the color-reversal task (rats) or across all tasks (tree shrews). Null distributions were constructed for spatial information content and for tuning curve MVLs (HD, FL, and EB). Cells were classified as significantly modulated by a given variable if their empirical values exceeded the 99^th^ percentile of the corresponding null distribution ^33^.

Slope differences: tetrode data and recording positions from both species were pooled and randomly resampled without replacement into two groups. For each iteration, the difference in regression slopes between the two groups was calculated. This process was repeated 10,000 times to generate a null distribution of slope differences. The observed slope difference was then compared against this distribution; the *P* value was defined as the proportion of permuted values exceeding the observed difference.

### Generalized additive model (GAM) fitting

Encoding of the six behavioral variables (position, HD, FL, EB, LS, and AV) was assessed using Poisson GAMs (neuroGAM; https://github.com/kaushik-l/neuroGAM) ^33^. Spike trains and behavioral variables were discretized into 20-ms bins; the spike-count vector was smoothed with a Gaussian kernel (σ = 3 bins). For each cell, the predicted firing rate *r* over *T* time points was modeled as:

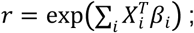

where *X_i_* represents the animal’s state for variable *i* at an instant of time, *β_i_* is the tuning curve for that variable.

Parameters were optimized by maximizing the log-likelihood (LLH) using MATLAB’s fminunc function (trust-region algorithm). The LLH *l* of observed spike train *n_t_* given predicted rate *r* across time points *t* was:

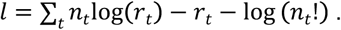

To prevent overfitting, a smoothing penalty *P*, was added to the objective function, penalizing differences between adjacent bins of each parameter vector:

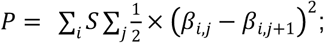

where *S* is a smoothing hyperparameter (set to 10 for all variables), *i* indexes over variables, and *j* indexes over response parameters for a given variable. Discretization: position (20 × 20 bins), FL (40 bins), HD and EB (36 bins each), LS and AV (10 bins each).

### Model validation and selection

Model significance was assessed using 10-fold cross-validation, and model performance was quantified as the LLH of held-out test data under the fitted model. Model significance was assessed using 10-fold cross-validation. Model selection followed an optimized forward procedure: (1) fit all single-variable models and retain the best if it outperformed a constant-rate null; (2) iteratively add variables until no additional predictor significantly improved LLH. The *dominant variable* for each neuron was defined as the predictor from the best-performing first-order (single-variable) model. The *complete encoding profile* was defined by variables included in the final best-fitting multivariable model.

Model significance was assessed using 10-fold cross-validation with 90% of the data used for training and 10% for testing. Significance relative to a constant-rate null model was determined using a one-sided Wilcoxon signed-rank test (P < 0.05). Model selection followed an optimized forward procedure: all single-variable models were first fitted; if any significantly outperformed the null model, the best-performing predictor was retained. That predictor was then combined with each remaining variable to fit all possible two-variable models, and the best additional variable was retained if it significantly improved fit. Variables were added sequentially until no additional predictor significantly improved LLH. The final model was the most parsimonious model achieving the highest significant LLH, favoring simpler models when addition of a variable did not yield a significant improvement.

### Multi-session neural data-sharing decoder

Classical Bayesian decoders require large within-session populations. Because wireless tree shrew recordings often yielded small populations, we adopted a multi-session data-sharing framework ^37^ (https://github.com/yzhang511/neural_decoding). The decoder predicted the animal’s instantaneous position from recent CA1 population activity while accommodating sessions with different numbers and identities of neurons. Only experiments with ≥ 4 simultaneously recorded neurons were included.

Spike counts were binned in non-overlapping 20-ms bins. Let *S_i_ ɛ R^N_i_xT_i_^* denote the spike-count matrix for session *i*, where *Ni* is the number of neurons and *T_i_* is the number of time bins. To exploit temporally structured spiking, *S_i_* was divided into non-overlapping five-bin (100 ms) segments. The neural input for segment *k* was:

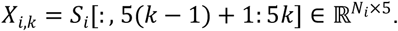

The behavioral target was the 2D position at the final bin of the segment:

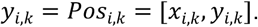

To share information across sessions without neuron identity matching, the decoder weight matrix for session *i* was factorized as:

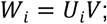

where *U_i_ ɛ R^N_i_xR^* maps session *i* neurons into an *R*-dimensional latent space, and *V ɛ R^RxT^* is shared across sessions. *U_i_* captured session-specific differences in recorded neuronal populations, whereas *V* captured temporal patterns consistently related to spatial behavior. For each segment, the model generated a prediction from the vectorized spike-count matrix:

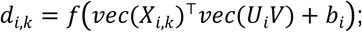

where *bi* is a session-specific bias term and *f* denotes the decoder output function. The position decoder was trained by minimizing the squared Euclidean loss:

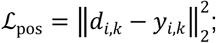

where *d_i,k_* and *y_i,k_* denote the network output and the ground-truth position at the final step of the *k*-th training sample, respectively.

For each species, data were pooled and randomly split into training (60%), validation (20%), and test (20%) sets. Model parameters were optimized on the training set, the best model was selected according to validation loss, and final performance was evaluated once on the test set. Tree shrew and rat models were trained independently with identical preprocessing, segmentation, architecture, optimization, and model-selection procedures. Decoding performance was quantified as the Euclidean position error on held-out test data. This approach allowed us to compare spatial localization performance between species employing different encoding strategies.

To control for potential sampling bias, rat sessions were rank-ordered by simultaneously recorded neuron count, and the 213 lowest-count sessions were selected to match the tree shrew dataset, yielding comparable neuron counts per session. This selection tested whether smaller tree shrew session sizes reflected a sampling ceiling; we confirmed that these low-count rat sessions were distributed across multiple animals and recording days. Decoding models were retrained on this matched dataset using identical procedures.

### Circular graph for conjunctive coding

Conjunctive coding was visualized and quantified using network graphs. Node size represents the fraction of neurons tuned to each variable. Line strength between variable pairs was quantified by the Jaccard index:

Graph layouts were generated using modified code from the Circular Graph toolbox (https://github.com/paul-kassebaum-mathworks/circularGraph).

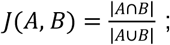

where *A* and *B* represent sets of neurons significantly tuned to variables *A* and *B*, respectively. Lines were rendered only for pairs with *J* > 0.1, with thickness proportional to *J* and color indicating the conditional proportion (fraction of neurons tuned to variable *A* that are also tuned to variable *B*). Graph layouts were generated using modified code from the Circular Graph toolbox.

### Two-Room Remapping Analysis

For two-room analyses, neurons were stratified by spatial information content (SIc) into non-positional (≤ 0.5 bits/spike), low-positional (0.5–1.0 bits/spike), and high-positional (≥ 1.0 bits/spike) groups.

Neurons were sorted by SIc into non-positional (≤ 0.5 bits/spike), low-positional (0.5–1.0 bits/spike), and high-positional (≥ 1.0 bits/spike) groups. Spatial correlation (SC) between rate maps (20 × 20 bins, Gaussian-smoothed) was computed via Pearson correlation between corresponding bins. Comparisons between inactive trials (mean firing rate < 0.1 Hz) were excluded. Rate maps from the second room were rotated in 90° increments to identify the alignment yielding maximum correlation.

Normalized changes in spatial tuning metrics across two trials were quantified using the following formula:

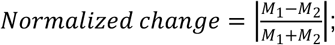

where *M_1_* and *M_2_* represent the compared parameter in trials 1 and 2, respectively.

Overall rate difference (RD), a metric for changes in overall activity level, was calculated as the normalized change in mean firing rate (Hz) between two trials.

Population vector correlation (PVC) was calculated by stacking individual rate maps into a 3D array (space × space × neurons). At each spatial bin, the population vector was defined as the vector of firing rates across all neurons. PVC between trials was the Pearson correlation between corresponding population vectors, using the rotational alignment that yielded the highest mean single-cell SC.

Preferred HD for each neuron was computed as the circular mean of its tuning curve in each room. Raw direction shifts across rooms were normalized by subtracting the animal-level mean shift. Normalized shifts were pooled and tested for directional concentration using the Rayleigh test. Pairwise angular differences between simultaneously recorded HD neuron pairs were computed per experiment and averaged for Moore’s paired-sample circular test.

### Statistics and reproducibility

Data are reported as mean ± standard error of the mean (SEM), or medians and interquartile ranges (IQR), depending on normality. One-sample Kolmogorov-Smirnov tests and Wilcoxon signed-rank tests examined uniformity and deviations above null distributions, respectively. Pairwise comparisons used Student’s *t*-test or Wilcoxon rank-sum/signed-rank tests as appropriate. Group comparisons used Friedman tests. Cross-species comparisons used two-way rank-transformed ANOVA (RT-ANOVA), with subregion and species as between-subject factors. PVCs were analyzed using repeated-measures two-way RT-ANOVA (subregion × group, both as within-subject factors). Linear mixed-effects models examined the fixed effects of species, CA1 subregions, and their interaction on spatial coding metrics; with random intercepts for neurons nested within animals. Categorical variables used Pearson’s chi-squared test. Linear associations used Pearson correlation coefficients. Significant spike-phase coupling was assessed per neuron using the Rayleigh test. *Post-hoc* pairwise comparisons used the Holm-Bonferroni method. Multiple comparisons between LFP frequency bands used false discovery rate (FDR) correction.

All tests were two-tailed with *P* < 0.05 considered significant, except for Rayleigh tests and shuffle-based permutation analyses for spatial coding (*P* < 0.01) ^5^ and GAM-related analyses (one-sided tests). Statistical analyses were performed using GraphPad Prism (v9; GraphPad Software, USA; RRID: SCR_002798) and figures generated using Origin (v10.1; OriginLab Corporation, USA; RRID: SCR_002815).

## Data Availability

Tree shrew single-cell transcriptomic data are deposited in NGDC GSA (CRA043801; to be released upon publication of the companion study). Other data supporting the findings of this study are provided in the Supplementary Data files. Additional datasets are available from the corresponding author upon reasonable request.

## Code Availability

Custom analysis code is available as Supplementary Code. Standard algorithms and publicly available software tools used in the analyses are referenced in the Methods section.

