## Supplementary Information for "Gradual evolution of multiplexed spatial coding within a conserved hippocampal scaffold"

**Extended Data Fig. 1 | Tree shrew CA1 recording sites**

**Extended Data Fig. 2 | Representative spike sorting examples**

**Extended Data Fig. 3 | Position-related firing properties in both species**

**Extended Data Fig. 4 | Tuning of spatially modulated neurons by permutation testing**

**Extended Data Fig. 5 | GAM fitting and coding strength by log-likelihood increase**

**Extended Data Fig. 6 | Species differences but subregional similarity in spatial variable coding**

**Extended Data Fig. 7 | Extended analyses for the light-dark task**

**Extended Data Fig. 8 | Extended analyses for the two-room task**

**Extended Data Fig. 9 | Gradual rodent-to-primate transition in hippocampal spatial coding**

**Supplementary Table 1 | Statistical analyses and outcomes**

**Supplementary Table 2 | Comparison of two-way RT-ANOVA and linear mixed-effects models**

**Supplementary Video 1 | Tree shrew open-field navigation**

### Extended Data Fig. 1

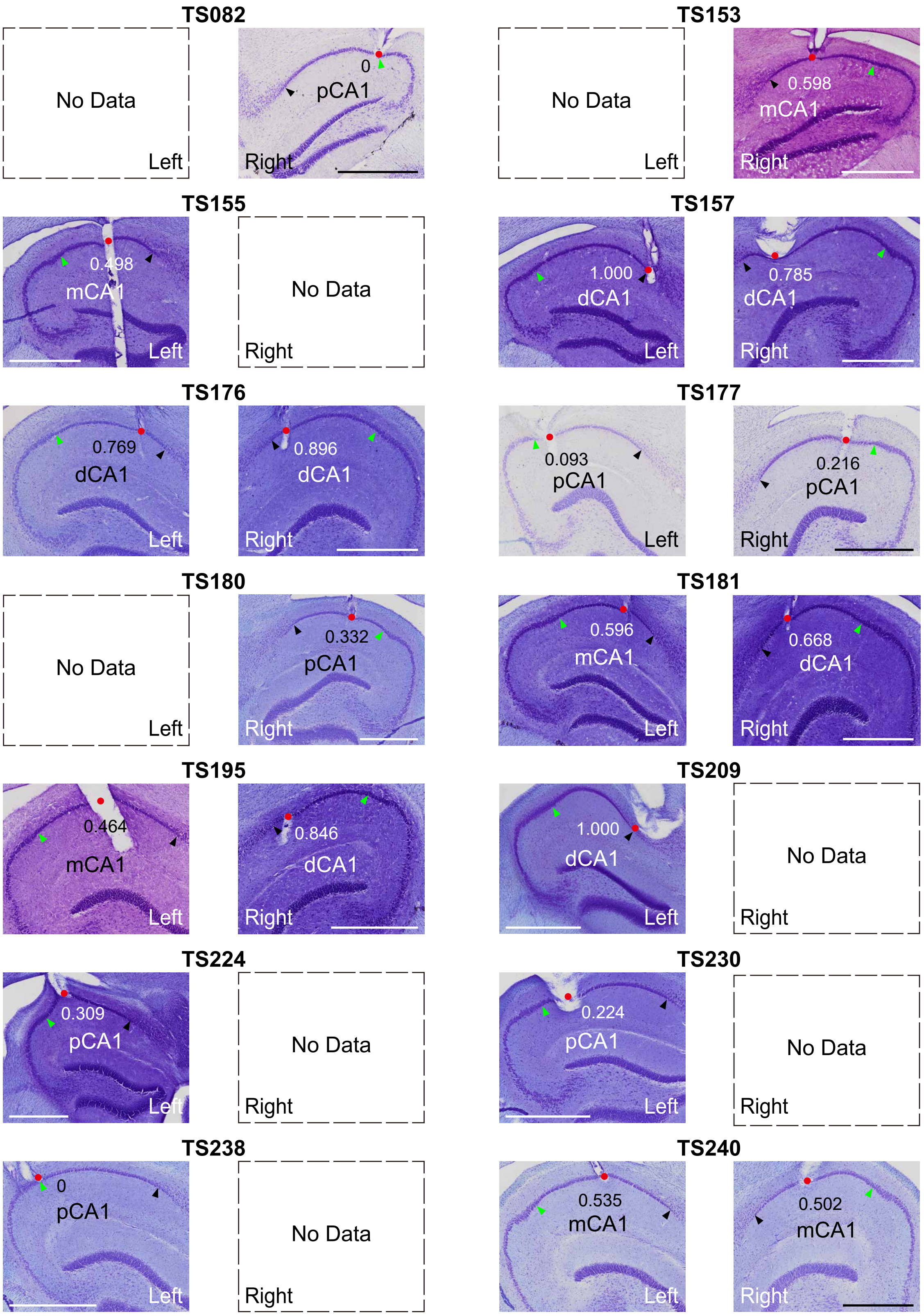


**Extended Data Fig. 1 | Tree shrew CA1 recording sites.**

Nissl-stained sagittal sections showing CA1 recording sites (red dots) for each animal. Numbers, normalized positions. Black/green arrowheads, CA1/subiculum and CA2/CA1 borders, respectively. For TS155 and TS195, tetrodes extended deeper, but data were collected at ~3500 µm. Scale: 1 mm.

### Extended Data Fig. 2

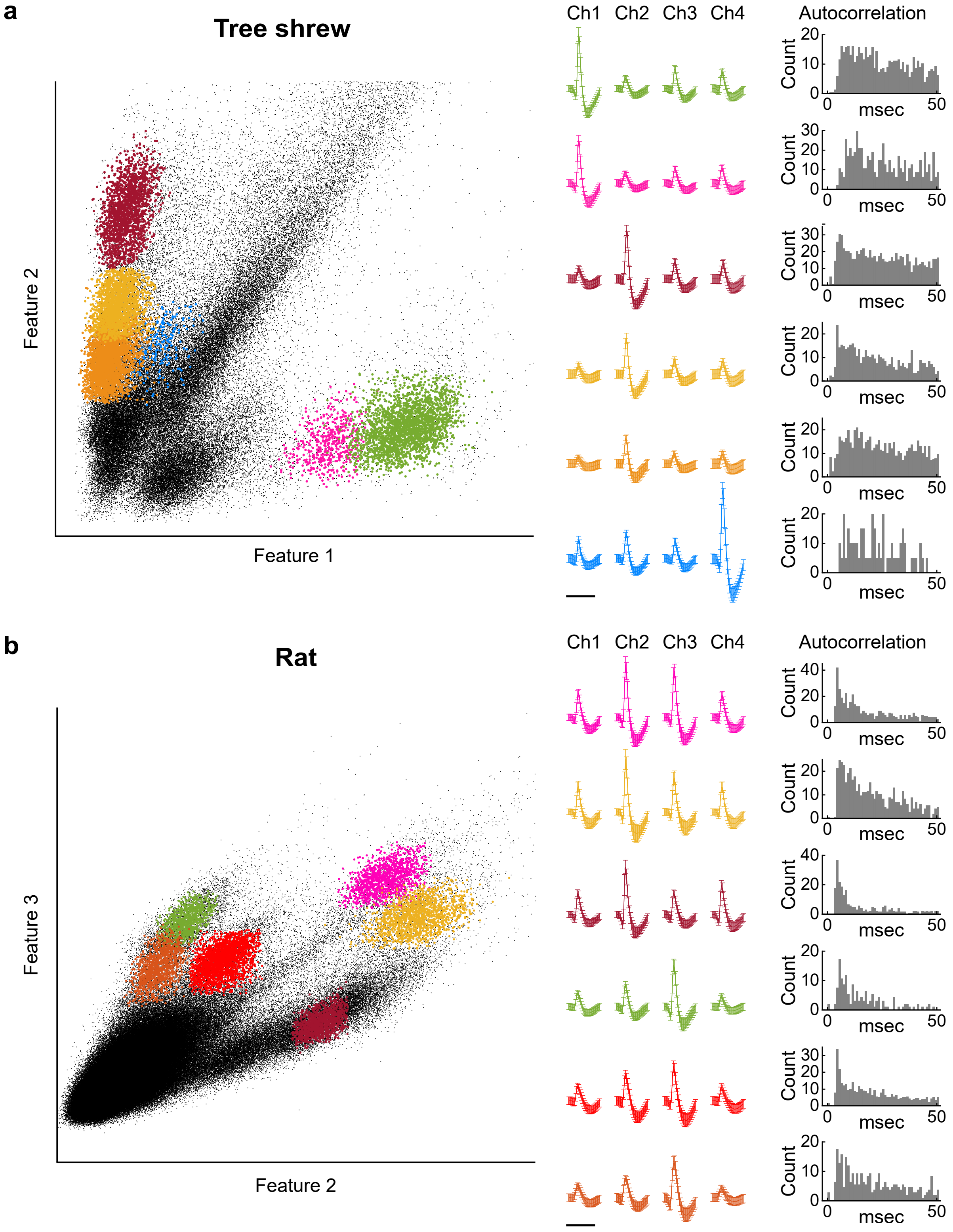


**Extended Data Fig. 2 | Representative spike sorting examples.**

**a**,Tree shrew CA1 tetrode. Left: 2D cluster diagrams (dots, spikes; clusters, putative single neurons; six color-coded clusters). Middle: mean waveforms (± SD). Right: autocorrelation histograms.

**b**,Rat CA1 tetrode, as in **a**. Scale: 1 ms.

### Extended Data Fig. 3

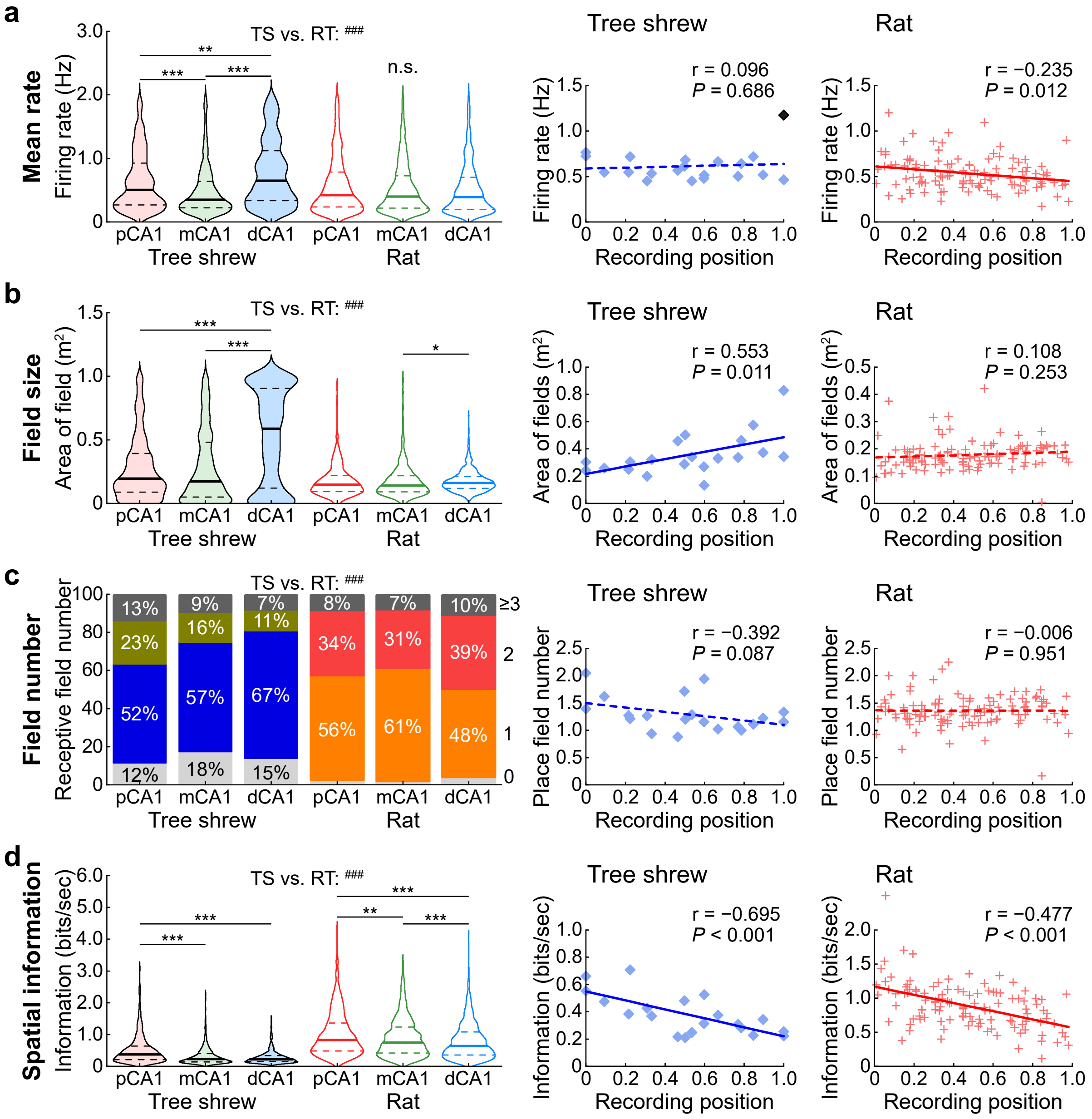

**Extended Data Fig. 3 | Position-related firing properties in both species.**

**a**, Mean firing rates (median (IQR); Hz). Tree shrews (TS), pCA1, 0.506 (0.265–0.926), mCA1, 0.349 (0.225–0.642), dCA1, 0.649 (0.333–1.124) (422, 483 and 420 neurons, respectively); rats (RT), pCA1, 0.422 (0.236–0.785), mCA1, 0.401 (0.217–0.726), dCA1, 0.389 (0.197–0.704) (863, 940 and 773 neurons). Two-way rank-transformed ANOVA (RT-ANOVA): species, $F(1, 3895) = 35.547$, $P < 0.001$, $\eta^2 = 0.009$; subregion, $F(2, 3895) = 24.081$, $P < 0.001$, $\eta^2 = 0.012$; species × subregion, $F(2, 3895) = 30.327$, $P < 0.001$, $\eta^2 = 0.015$. Right: rates vs. normalized CA1 transverse axis (20 TS sites; 113 RT sites); solid line, significant regression; dashed, non-significant. No slope difference (permutation test, $P = 0.136$). Dark blue dot, tetrodes from TS209 at the CA1–subiculum border potentially capturing subicular activity. Exclusion of this site abolished the elevated mean rate in dCA1 but did not alter dCA1 spatial coding metrics.

**b**–**d**, Position-related metrics, displayed as in **a**. **b**, Receptive field size m$^2$: TS, pCA1, 0.196 (0.088–0.392), mCA1, 0.173 (0.050–0.480), dCA1, 0.588 (0.119–0.907); RT, pCA1, 0.148 (0.093–0.220), mCA1, 0.141 (0.090–0.219), dCA1, 0.160 (0.117–0.212). Species, $F(1, 3895) =143.664$, $P < 0.001$, $\eta^2 = 0.036$; subregion, $F(2, 3895) = 51.818$, $P < 0.001$, $\eta^2 = 0.026$; species × subregion, $F(2, 3895) = 21.608$, $P < 0.001$, $\eta^2 = 0.011$. Right: transverse gradient steeper in tree shrews ($P = 0.031$). **c**, Receptive field number: TS, pCA1, 1 (1–2), mCA1, 1 (1–1), dCA1, 1 (1–1); RT, pCA1, 1 (1–1.5), mCA1, 1 (1–1.5), dCA1, 1 (1–2). Species, $F(1, 3895) = 82.687$, $P < 0.001$, $\eta^2 = 0.021$; subregion, $F(2, 3895) = 11.121$, $P < 0.001$, $\eta^2 = 0.006$; species × subregion, $F(2, 3895) = 17.159$, $P < 0.001$, $\eta^2 = 0.009$. Right: no slope difference ($P = 0.073$). **d**, Spatial information rate (bits/second): TS, pCA1, 0.370 (0.207–0.641), mCA1, 0.226 (0.138–0.400), dCA1, 0.219 (0.140–0.332); RT, pCA1, 0.823 (0.477–1.358), mCA1, 0.747 (0.416–1.231), dCA1, 0.635 (0.351–1.078). Species, $F(1, 3895) = 1362.754$, $P < 0.001$, $\eta^2 = 0.259$; subregion, $F(2, 3895) = 83.272$, $P < 0.001$, $\eta^2 = 0.041$; species × subregion, $F(2, 3895) = 14.198$, $P < 0.001$, $\eta^2 = 0.007$. Right: no slope difference ($P = 0.219$).

n.s., not significant; cross-region: *$P < 0.05$, **$P < 0.01$, ***$P < 0.001$; cross-species: $^{\#\#\#}P < 0.001$.

### Extended Data Fig. 4

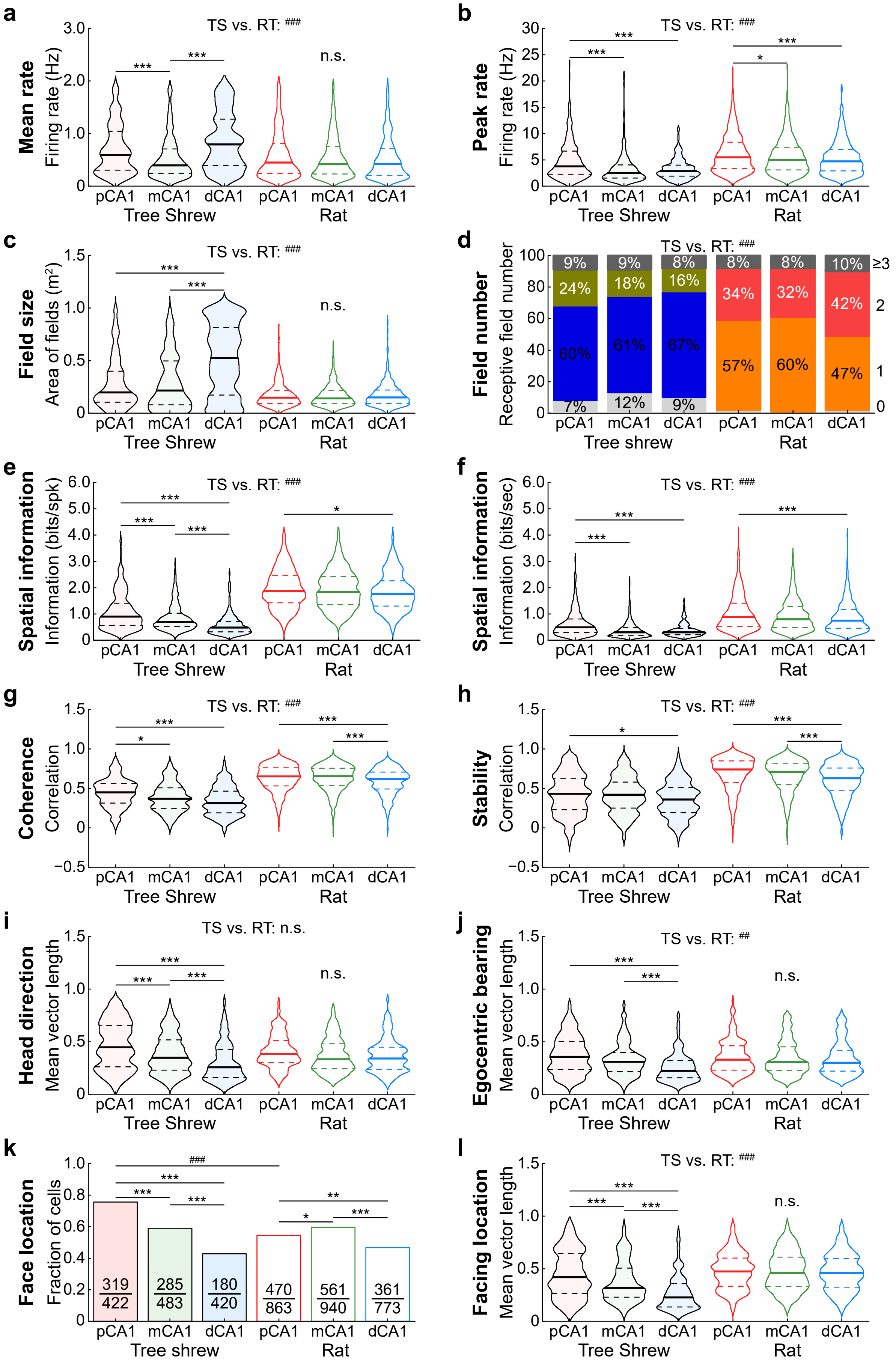

**Extended Data Fig. 4 | Tuning of spatially modulated neurons by permutation testing.**

**a**–**h**, Position-selective neurons (median (IQR)). Values for pCA1/mCA1/dCA1 are listed in order. **a**, Mean firing rate (Hz). TS: 0.591 (0.302–1.048) / 0.395 (0.244–0.710) / 0.795 (0.391–1.279) (n = 289/307/182); RT: 0.448 (0.244–0.814) / 0.417 (0.230–0.754) / 0.421 (0.204–0.719) (n = 817/878/675). Two-way RT-ANOVA, species, $F(1, 3142) = 55.626$, $P < 0.001$, $\eta^2 = 0.017$; subregion, $F(2, 3142) = 18.387$, $P < 0.001$, $\eta^2 = 0.012$; species × subregion, $F(2, 3142) = 20.218$, $P < 0.001$, $\eta^2 = 0.013$. **b**, Peak firing rate (Hz). TS: 3.774 (2.275–6.754) / 2.491 (1.526–4.042) / 2.850 (1.911–4.010); RT: 5.473 (3.353–8.342) / 5.009 (3.068–7.419) / 4.700 (2.911–7.010). Species, $F(1, 3142) = 291.523$, $P < 0.001$, $\eta^2 = 0.085$; subregion, $F(2, 3142) = 35.009$, $P < 0.001$, $\eta^2 = 0.022$; species × subregion, $F(2, 3142) = 9.753$, $P < 0.001$, $\eta^2 = 0.006$. **c**, Place field size ($m^2$). TS: 0.198 (0.105–0.401) / 0.215 (0.080–0.498) / 0.524 (0.173–0.818); RT: 0.148 (0.094–0.216) / 0.141 (0.091–0.215) / 0.149 (0.093–0.221). Species, $F(1, 3142) = 193.714$, $P < 0.001$, $\eta^2 = 0.058$; subregion, $F(2, 3142) = 23.782$, $P < 0.001$, $\eta^2 = 0.015$; species × subregion, $F(2, 3142) = 19.346$, $P < 0.001$, $\eta^2 = 0.012$. **d**, Place field number. TS: 1 (1–2) / 1 (1–2) / 1 (1–1); RT: 1 (1–1.5) / 1 (1–1.5) / 1.5 (1–2). Species, $F(1, 3142) = 39.615$, $P < 0.001$, $\eta^2 = 0.012$; subregion, $F(2, 3142) = 3.809$, $P = 0.022$, $\eta^2 = 0.002$; species × subregion, $F(2, 3142) = 7.831$, $P < 0.001$, $\eta^2 = 0.005$. **e**, Spatial information content (SIc; bits/spike). TS: 0.901 (0.555–1.417) / 0.698 (0.517–1.029) / 0.479 (0.318–0.709); RT: 1.868 (1.419–2.461) / 1.838 (1.352–2.424) / 1.763 (1.303–2.263). Species, $F(1, 3142) = 1486.849$, $P < 0.001$, $\eta^2 = 0.321$; subregion, $F(2, 3142) = 37.570$, $P < 0.001$, $\eta^2 = 0.023$; species × subregion, $F(2, 3142) = 15.281$, $P < 0.001$, $\eta^2 = 0.010$. **f**, Spatial information rate (bits/second). TS: 0.486 (0.295–0.805) / 0.300 (0.172–0.481) / 0.302 (0.212–0.445); RT: 0.877 (0.521–1.408) / 0.798 (0.481–1.282) / 0.742 (0.450–1.172). Species, $F(1, 3142) = 664.543$, $P < 0.001$, $\eta^2 = 0.175$; subregion, $F(2, 3142) = 43.456$, $P < 0.001$, $\eta^2 = 0.027$; species × subregion, $F(2, 3142) = 12.438$, $P < 0.001$, $\eta^2 = 0.008$. **g**, Spatial coherence. TS: 0.449 (0.313–0.563) / 0.369 (0.248–0.507) / 0.313 (0.188–0.465); RT: 0.652 (0.533–0.763) / 0.656 (0.539–0.757) / 0.620 (0.493–0.707). Species, $F(1, 3142) = 905.980$, $P < 0.001$, $\eta^2 = 0.224$; subregion, $F(2, 3142) = 24.105$, $P < 0.001$, $\eta^2 = 0.015$; species × subregion, $F(2, 3142) = 4.123$, $P = 0.016$, $\eta^2 = 0.003$. **h**, Spatial stability. TS: 0.432 (0.228–0.629) / 0.421 (0.252–0.580) / 0.360 (0.191–0.515); RT: 0.739 (0.572–0.850) / 0.710 (0.548–0.820) / 0.628 (0.472–0.759). Species, $F(1, 3138) = 678.055$, $P < 0.001$, $\eta^2 = 0.178$; subregion, $F(2, 3138) = 26.300$, $P < 0.001$, $\eta^2 = 0.016$; species × subregion, $F(2, 3138) = 1.340$, $P = 0.262$.

**i**–**l**, Non-positional tuning. **i**, Head direction (HD) mean vector length (MVL) for HD-modulated cells. TS: 0.446 (0.262–0.653) / 0.347 (0.227–0.516) / 0.257 (0.159–0.426) (n = 316/292/179); RT: 0.384 (0.301–0.515) / 0.333 (0.242–0.483) / 0.341 (0.235–0.448) (N = 248/202/140). Species, $F(1, 1371) = 1.793$, $P = 0.181$; subregion, $F(2, 1371) = 32.870$, $P < 0.001$, $\eta^2 = 0.046$; species × subregion, $F(2, 1371) = 9.229$, $P < 0.001$, $\eta^2 = 0.013$. **j**, Egocentric bearing (EB) MVL for EB-modulated cells. TS: 0.357 (0.235–0.501) / 0.310 (0.214–0.398) / 0.223 (0.155–0.321) (n = 203/218/186); RT: 0.328 (0.229–0.461) / 0.306 (0.226–0.452) / 0.300 (0.220–0.423) (n = 165/136/93). Species, $F(1, 995) = 7.460$, $P = 0.006$, $\eta^2 = 0.007$; subregion, $F(2, 995) = 15.327$, $P < 0.001$, $\eta^2 = 0.030$; species × subregion, $F(2, 995) = 8.660$, $P < 0.001$, $\eta^2 = 0.017$. *Post-hoc* TS pCA1 vs. RT pCA1, mCA1, and dCA1, all $P = 1.000$. **k**, Proportion of facing location (FL)-tuned neurons. TS vs. RT: pCA1, $\chi^2(1) = 53.40$, $P < 0.001$, $\varphi = 0.204$; mCA1, $\chi^2(1) = 0.060$, $P = 1.000$; dCA1, $\chi^2(1) = 1.623$, $P = 0.609$. Regional: TS, $\chi^2(2) = 93.38$, $P < 0.001$, Cramer's $V = 0.265$; RT, $\chi^2(2) = 28.87$, $P < 0.001$, Cramer's $V = 0.106$. **l**, FL MVL for FL-modulated cells. TS: 0.420 (0.265–0.644) / 0.318 (0.228–0.506) / 0.227 (0.137–0.360) (n = 319/285/180). RT: 0.474 (0.333–0.601) / 0.461 (0.330–0.614) / 0.460 (0.323–0.597) (n = 470/561/361). Species, $F(1, 2170) = 183.156$, $P < 0.001$, $\eta^2 = 0.078$; subregion, $F(2, 2170) = 36.444$, $P < 0.001$, $\eta^2 = 0.033$; species × subregion, $F(2, 2170) = 28.195$, $P < 0.001$, $\eta^2 = 0.025$. *Post-hoc* TS pCA1 vs. rat pCA1, mCA1, and dCA1, all $P > 0.058$.

n.s., not significant; cross-region: $^{*}P < 0.05$, $^{**}P < 0.01$, $^{***}P < 0.001$; cross-species: $^{\#\#}P < 0.01$, $^{\#\#\#}P < 0.001$.

**Extended Data Fig. 5**

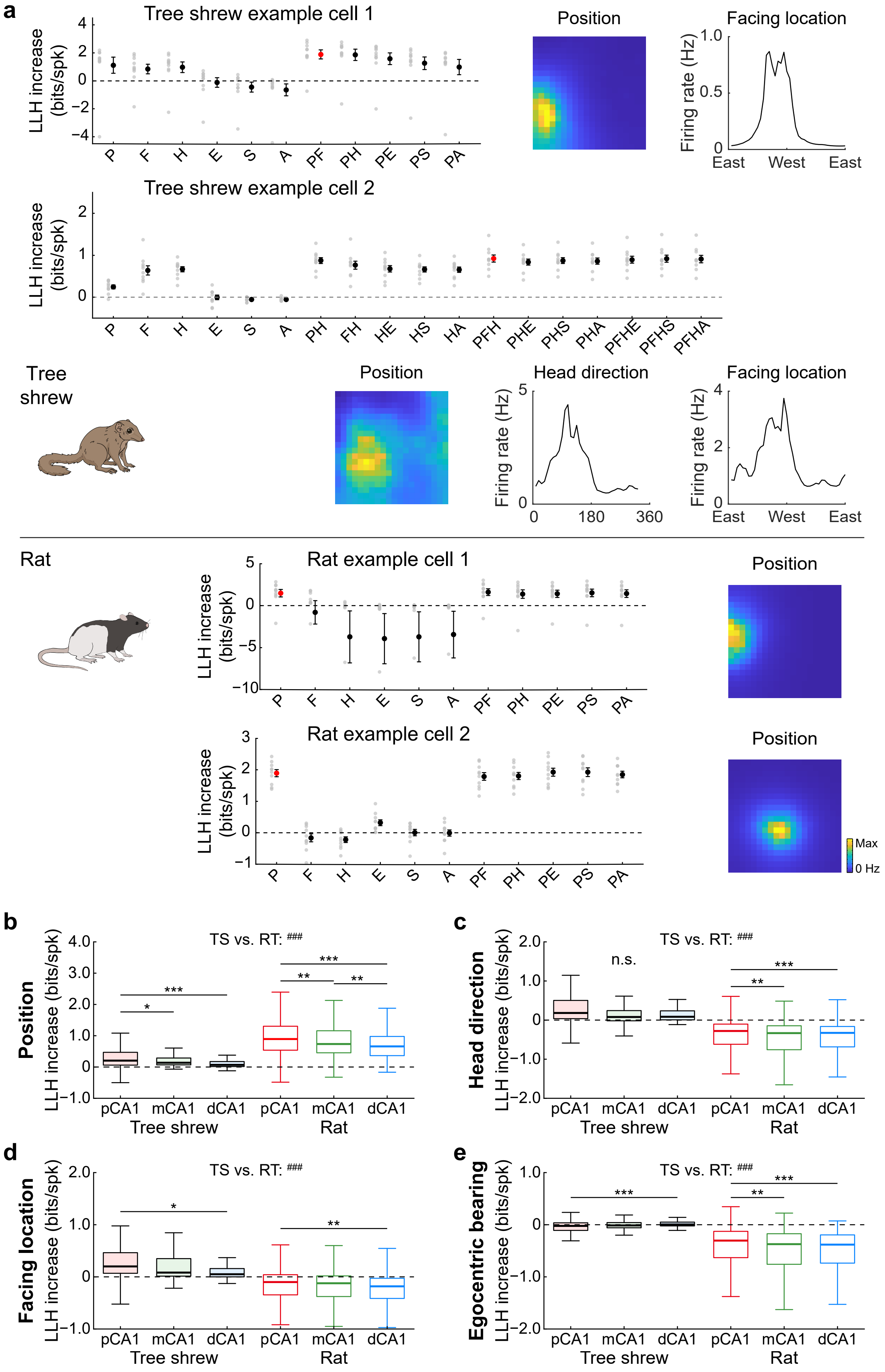

**Extended Data Fig. 5 | GAM fitting and coding strength by log-likelihood increase.**

**a**, Generalized additive model (GAM) fitting for Figure 4A examples (forward selection). Gray dots, log-likelihood (LLH) increase for each predictor added to the null model; black dots, mean ± SEM; red dot, best model. TS neuron 1: conjunctive position (P) and facing location (F); TS neuron 2: P, F, and head direction (H). Rat neurons: position only. E, egocentric bearing; S, linear speed; A, angular velocity.

**b**–**e**, Coding strength (LLH increase relative to null; bits/spike; median (IQR)). Values for pCA1/mCA1/dCA1 in order. TS sample sizes after removing NaNs: 221/205/120; RT: 583/588/395. **b**, Position. TS: 0.204 (0.060–0.476) / 0.139 (0.067–0.289) / 0.066 (0.015–0.171). RT: 0.895 (0.539–1.306) / 0.734 (0.457–1.162) / 0.662 (0.365–0.975). Two-way RT-ANOVA: species, $F(1, 2106) = 854.948$, $P < 0.001$, $\eta^2 = 0.289$; subregion, $F(2, 2106) = 34.249$, $P < 0.001$, $\eta^2 = 0.032$; species × subregion, $F(2, 2106) = 0.859$, $P = 0.424$. RT exceeded TS in all subregions (all $P < 0.001$). All LLH > 0, one-sided one-sample Wilcoxon signed-rank test against zero, TS: pCA1, Z = 11.358, $P < 0.001$, $r = 0.764$; mCA1, Z = 11.713, $P < 0.001$, $r = 0.818$; dCA1, Z = 7.483, $P < 0.001$, $r = 0.683$; RT: pCA1, Z = 20.678, $P < 0.001$, $r = 0.856$; mCA1, Z = 20.433, $P < 0.001$, $r = 0.843$; dCA1, Z = 16.944, $P < 0.001$, $r = 0.853$. **c**, HD. TS: 0.179 (0.038–0.504) / 0.078 (−0.020–0.249) / 0.084 (0.009–0.238). RT: −0.279 (−0.615–−0.101) / −0.333 (−0.758–−0.146) / −0.329 (−0.683–−0.164). Species, $F(1, 2106) = 1593.863$, $P < 0.001$, $\eta^2 = 0.431$; subregion, $F(2, 2106) = 6.545$, $P = 0.001$, $\eta^2 = 0.006$; species × subregion, $F(2, 2106) = 1.150$, $P = 0.317$. LLH > 0 in TS (pCA1, Z = 9.947, $P < 0.001$, $r = 0.669$; mCA1, Z = 7.424, $P < 0.001$, $r = 0.519$; dCA1, Z = 7.376, $P < 0.001$, $r = 0.673$), but not in RT (all Z < −16.7, all $P = 1.000$). **d**, FL. TS: 0.202 (0.065–0.470) / 0.084 (0.012–0.351) / 0.051 (0.001–0.163). RT: −0.100 (−0.346–0.042) / −0.123 (−0.376–0.019) / −0.184 (−0.414–−0.030). Species, $F(1, 2106) = 650.552$, $P < 0.001$, $\eta^2 = 0.236$; subregion, $F(2, 2106) = 11.959$, $P < 0.001$, $\eta^2 = 0.011$; species × subregion, $F(2, 2106) = 1.188$, $P = 0.305$. LLH > 0 in TS (pCA1, Z = 11.535, $P < 0.001$, $r = 0.776$; mCA1, Z = 9.440, $P < 0.001$, $r = 0.659$; dCA1, Z = 6.635, $P < 0.001$, $r = 0.606$), but not in RT (all Z < −9.7, all $P = 1.000$). **e**, EB. TS: −0.023 (−0.109–0.037) / −0.016 (−0.059–0.040) / 0.003 (−0.022–0.049). RT: −0.305 (−0.630–−0.128) / −0.372 (−0.759–−0.172) / −0.380 (−0.738–−0.195). Species, $F(1, 2106) = 1237.615$, $P < 0.001$, $\eta^2 = 0.370$; subregion, $F(2, 2106) = 1.279$, $P = 0.278$; species × subregion, $F(2, 2106) = 16.817$, $P < 0.001$, $\eta^2 = 0.016$. LLH not >0 in either species (TS: pCA1, Z = −4.071, $P = 1.000$; mCA1, Z = −2.208, $P = 0.986$; dCA1, Z = 1.531, $P = 0.063$; RT: all Z < −17.1, all $P = 1.000$).

n.s., not significant; cross-region: *$P < 0.05$, **$P < 0.01$, ***$P < 0.001$; cross-species: $^{\#\#\#}P < 0.001$.

**Extended Data Fig. 6-1**

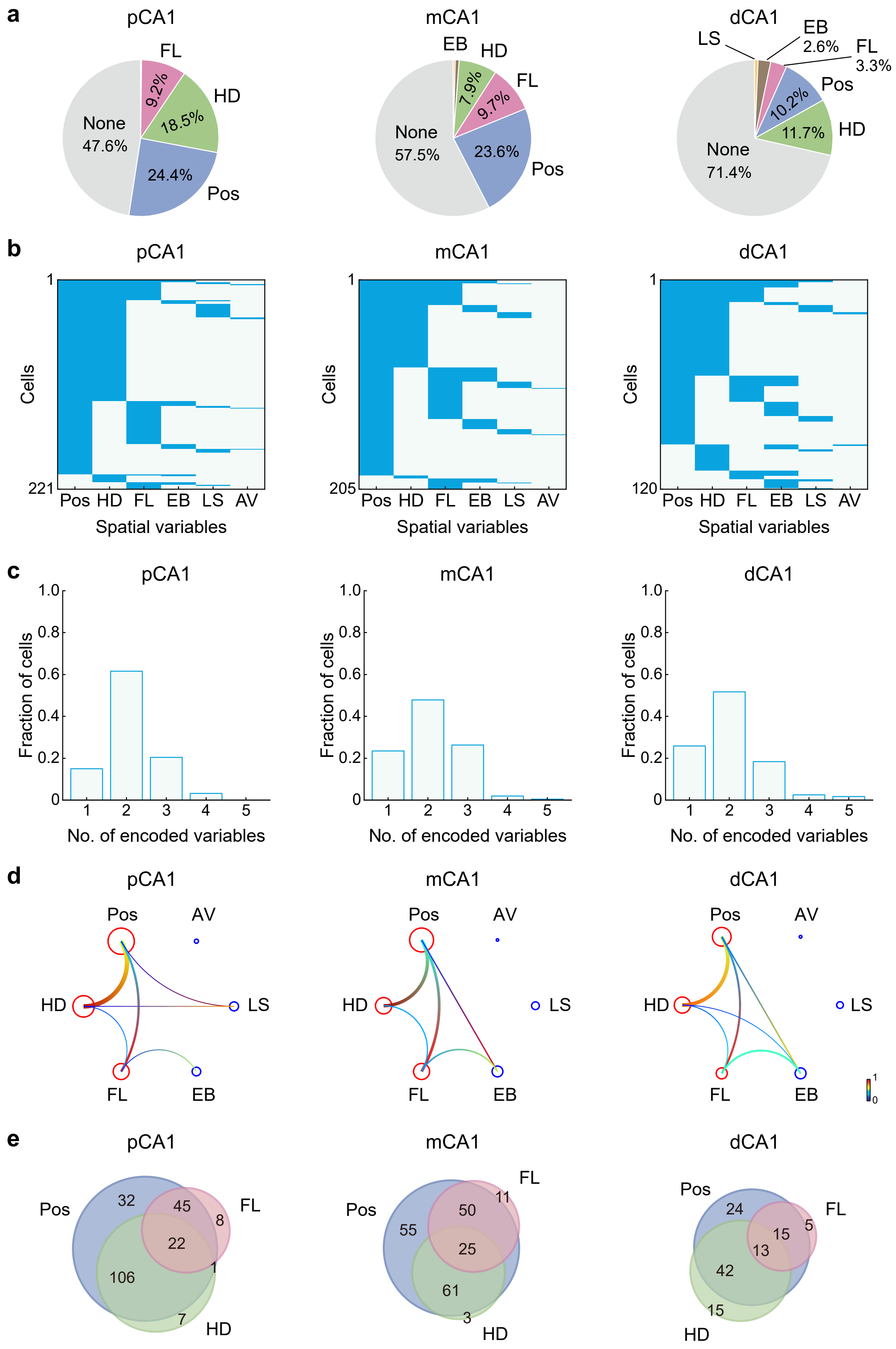

**Extended Data Fig. 6-2**

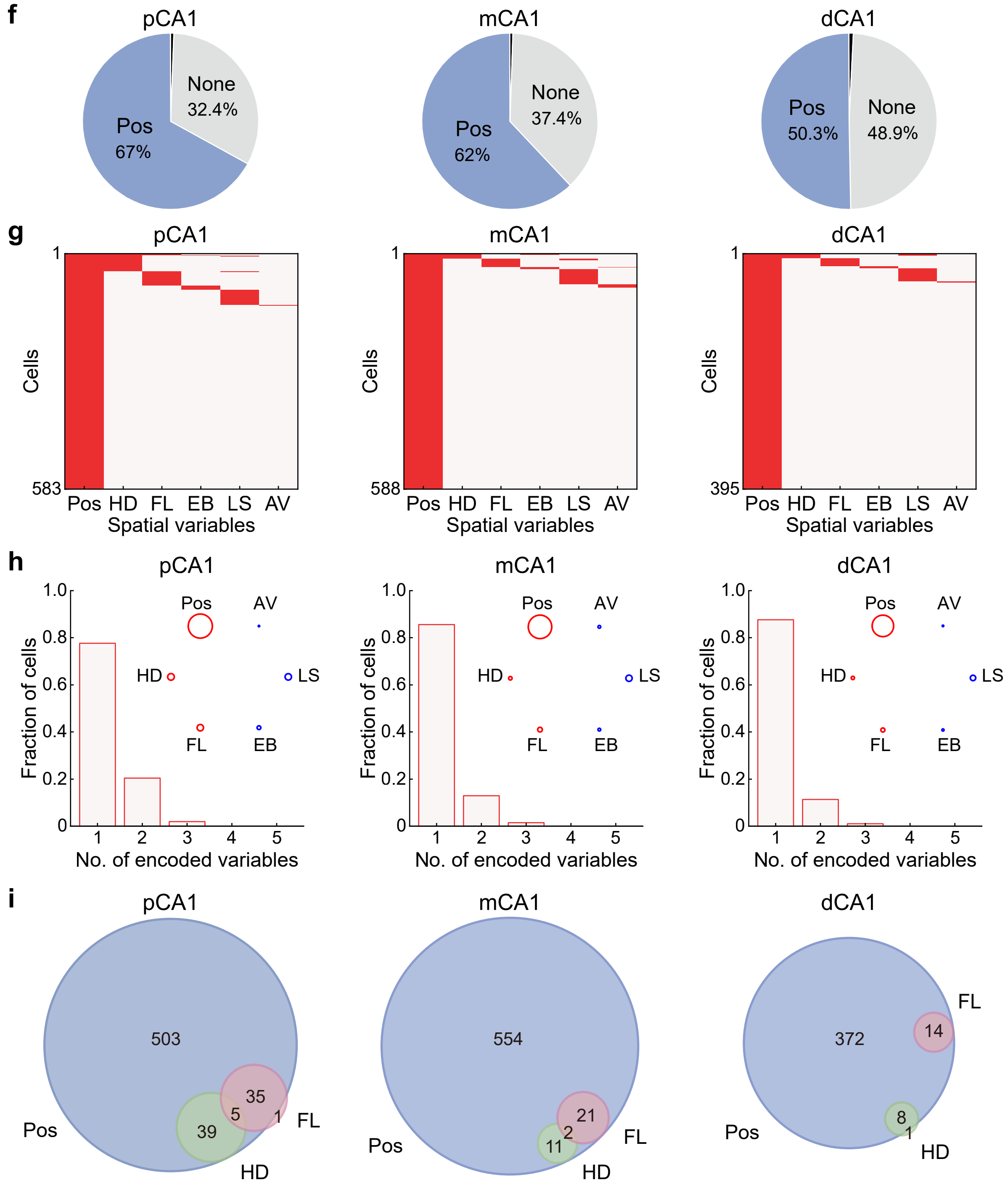


**Extended Data Fig. 6 | Species differences but subregional similarity in spatial variable coding.**

**a**–**e**, Tree shrew CA1 subregional coding. **a**, Pie charts: proportions by dominant variable (Pos, position; LS, linear speed; AV, angular velocity). **b**, Variable combination matrix (columns: variables; rows: neurons; filled cells: significant tuning). **c**, Fraction encoding 1, 2, or 3+ variables. **d**, Circular graphs of conjunctive coding (node size, proportion; line thickness, Jaccard index; color, conditional proportion). Allocentric connections (Pos, HD, FL) stronger than others: Wilcoxon rank-sum test, pCA1, $Z = 2.382$, $P = 0.017$, $r = 0.615$; mCA1, $Z = 2.245$, $P = 0.025$, $r = 0.580$; dCA1, $Z = 2.239$, $P = 0.025$, $r = 0.578$. **e**, Venn diagrams of allocentric tuning overlap (numbers, neuron counts).

**f**–**i**, Rat CA1, as in **a**–**e**.

### Extended Data Fig. 7

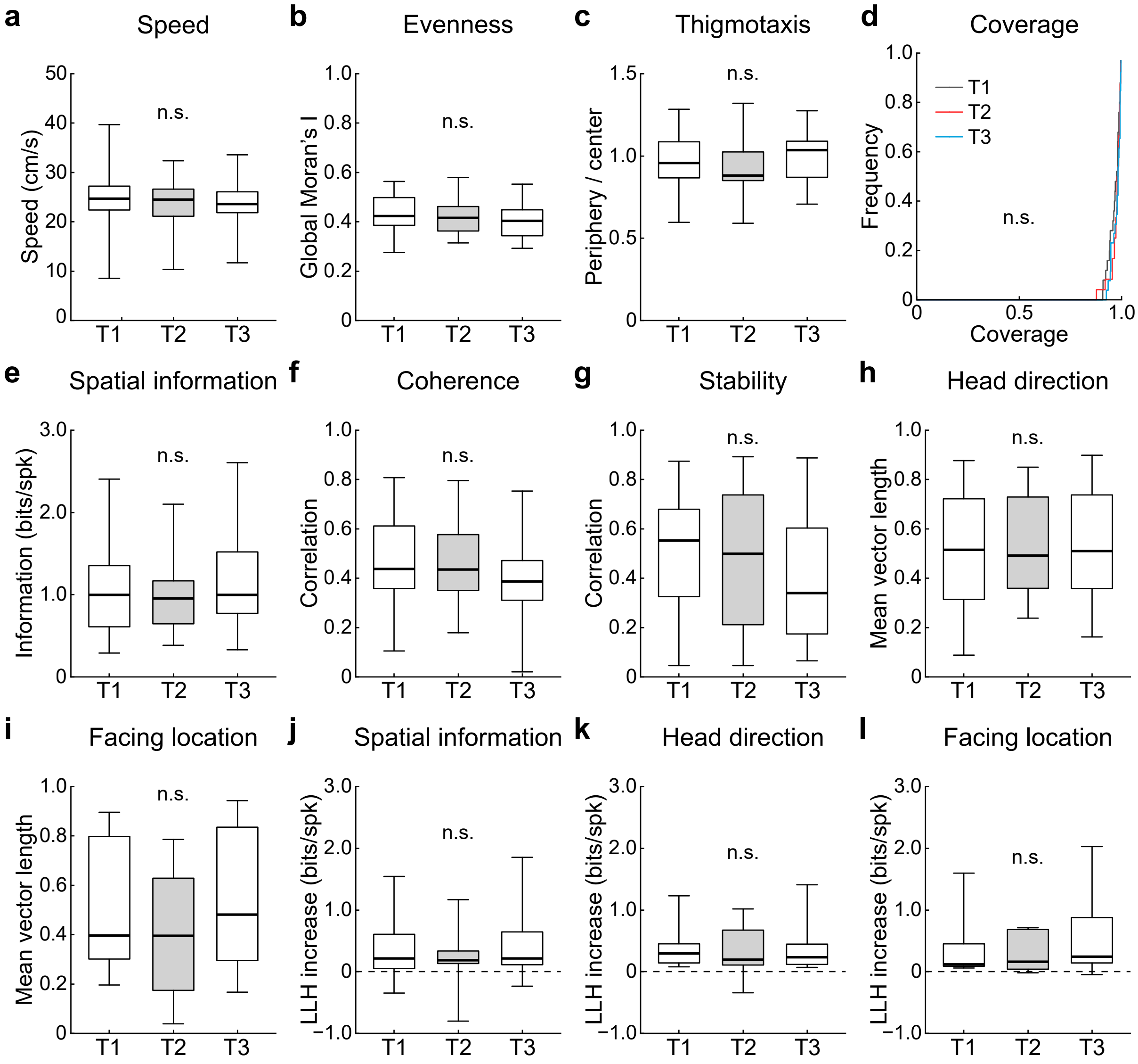

**Extended Data Fig. 7 | Extended analyses for the light-dark task.**

**a**–**d**, Behavioral and spatial sampling remained stable across lighting conditions (26 experiments; median (IQR)). **a**, Speed (cm/s). Trial 1: 24.796 (22.430–27.559); trial 2: 24.615 (21.111–26.970); trial 3: 23.709 (21.841–26.359); Friedman test, $\chi^2(2) = 1.231$, $P = 0.540$. **b**, Trajectory evenness (Global Moran's I). Trial 1: 0.424 (0.384–0.503); trial 2: 0.416 (0.363–0.463); trial 3: 0.404 (0.342–0.451); $\chi^2(2) = 1.000$, $P = 0.607$. **c**, Thigmotaxis. 1: 0.957 (0.866–1.089); trial 2: 0.882 (0.850–1.030); trial 3: 1.036 (0.869–1.098); $\chi^2(2) = 5.154$, $P = 0.076$. **d**, Arena coverage. Trial 1: 0.978 (0.944–0.991); trial 2: 0.983 (0.971–0.992); trial 3: 0.983 (0.959–0.993); $\chi^2(2) = 4.288$, $P = 0.117$.

**e**–**i**, Raw tuning metrics across trials (median (IQR)). **e**, SIc (bits/spike; n = 25). Trial 1: 0.996 (0.589–1.484); trial 2: 0.953 (0.645–1.167); trial 3: 0.999 (0.762–1.539); Mixed-effects analysis, $F(1.771, 40.73) = 1.060$, $P = 0.349$. **f**, Coherence. Trial 1: 0.438 (0.350–0.612); trial 2: 0.435 (0.350–0.576); trial 3: 0.387 (0.302–0.494); $F(1.836, 42.23) = 1.492$, $P = 0.237$. **g**, Stability. Trial 1: 0.552 (0.294–0.680); trial 2: 0.499 (0.212–0.737); trial 3: 0.339 (0.162–0.605); $F(1.917, 44.09) = 2.859$, $P = 0.070$. **h**, HD (n = 19). Trial 1: 0.515 (0.315–0.722); trial 2: 0.492 (0.345–0.732); trial 3: 0.511 (0.359–0.738); $F(1.967, 34.43) = 0.224$, $P = 0.797$. **i**, FL (n = 10). Trial 1: 0.397 (0.299–0.812); trial 2: 0.395 (0.142–0.673); trial 3: 0.481 (0.291–0.850); $F(1.345, 10.76) = 0.371$, $P = 0.618$.

**j**–**l**, LLH increase (bits/spike). **j**, Position. Trial 1: 0.204 (0.046–0.610); trial 2: 0.193 (0.126–0.347); trial 3: 0.215 (0.060–0.719); $F(1.531, 22.97) = 0.208$, $P = 0.755$. **k**, HD. Trial 1: 0.245 (0.142–0.453); trial 2: 0.207 (0.107–0.661); trial 3: 0.233 (0.111–0.500); $F(1.278, 15.98) = 0.075$, $P = 0.846$. **l**, FL. Trial 1: 0.116 (0.078–0.583); trial 2: 0.160 (0.040–0.682); trial 3: 0.242 (0.142–0.875); $F(0.513, 2.821) = 0.757$, $P = 0.352$.

n.s., not significant.

**Extended Data Fig. 8**

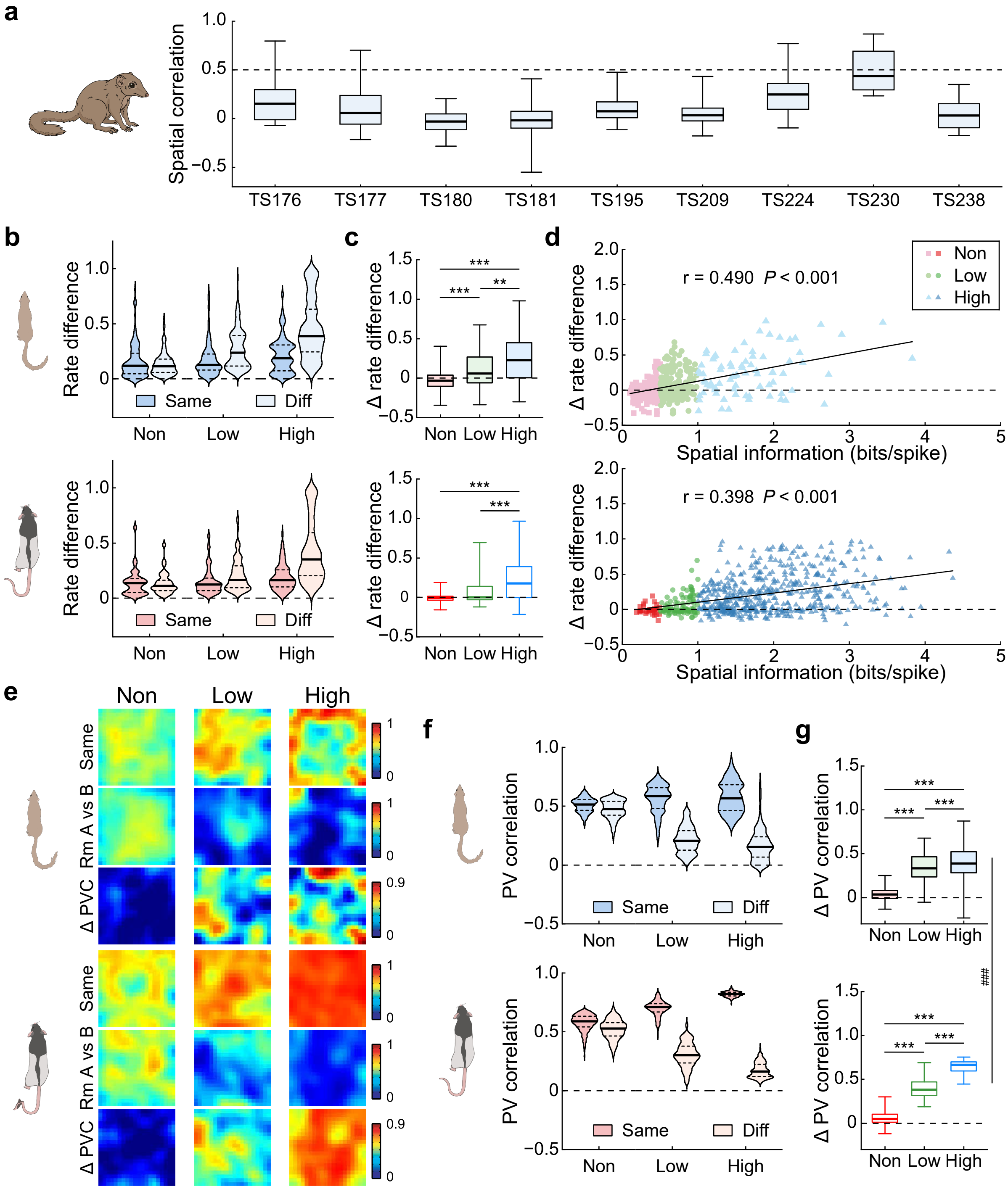

**Extended Data Fig. 8 | Extended analyses for the two-room task.**

**a**, Between-room spatial correlation of position-selective neurons (SIc > 0.5 bits/spike) for each tree shrew. All animals showed median correlations < 0.5, indicating global remapping.

**b**, Rate difference (RD) distributions for low-, middle-, and high-SIc groups. Dark blue/red: same-room; light blue/red: different-room. Solid line, median; dashed, quartiles.

**c**, Room-switch ΔRD by SIc group (outliers hidden). Two-way RT-ANOVA: species, $F(1, 993) = 0.003$, $P = 0.959$; neuron group, $F(2, 993) = 52.976$, $P < 0.001$, $\eta^2 = 0.096$; species × group interaction, $F(2, 993) = 0.810$, $P = 0.445$.

**d**, SIc vs. ΔRD. Tree shrews (top): non-positional, 155; low-positional, 148; high-positional, 79 neurons. Rats (bottom): non, 34; low, 91; high, 492. Solid line, significant regression; Pearson *r* and *P* indicated.

**e**, Population vector correlation (PVC) heatmaps sorted by SIc. Top: within-room; middle: across-room; bottom: ΔPVC.

**f**, PVC distributions across CA1 groups, as in **b**.

**g**, Room-switch ΔPVC. Repeated-measures two-way RT-ANOVA: species, $F(1, 2394) = 431.295$, $P < 0.001$, $\eta^2 = 0.153$; neuron group, $F(2, 2394) = 3419.294$, $P < 0.001$, $\eta^2 = 0.741$; species × group interaction, $F(2, 2394) = 184.869$, $P < 0.001$, $\eta^2 = 0.134$.

Cross-group: $^{**}P < 0.01$, $^{***}P < 0.001$; cross-species: $^{\#\#\#}P < 0.001$.

**Extended Data Fig. 9**

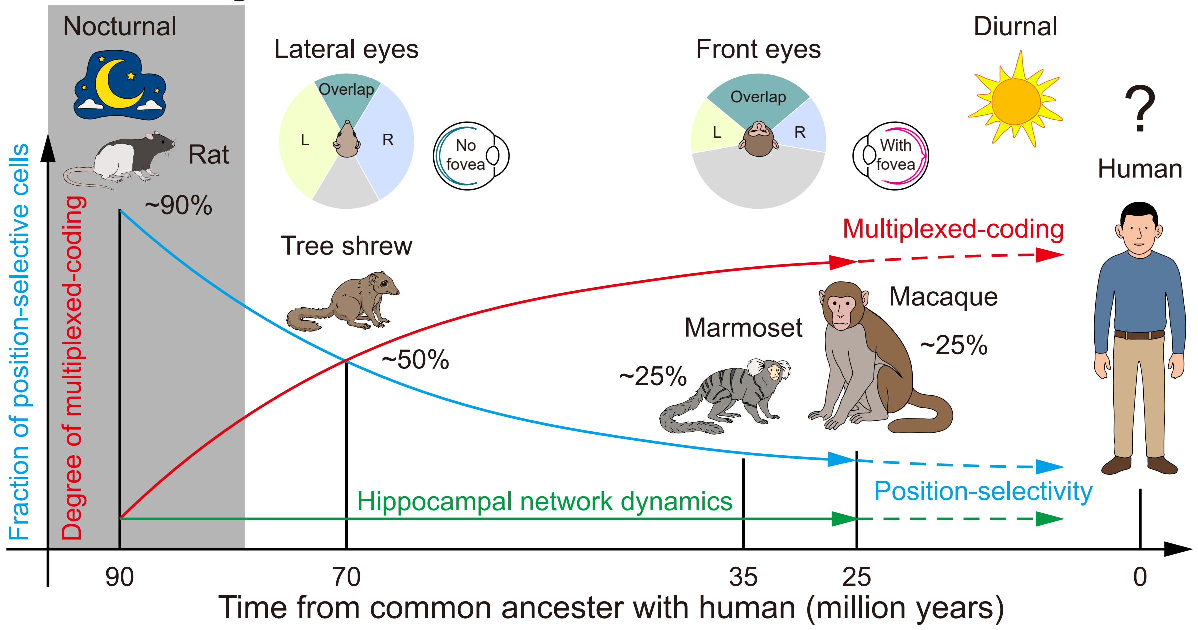


**Extended Data Fig. 9 | Gradual rodent-to-primate transition in hippocampal spatial coding.** Schematic of hippocampal spatial coding evolution across mammals. Rodents show predominantly position-dominant representations; primates exhibit weaker position modulation with prevalent multiplexed tuning to multiple spatial variables. Tree shrews, as a phylogenetically intermediate species, display a hybrid profile bridging these extremes: attenuated position selectivity relative to rodents alongside enhanced non-positional and multiplexed coding approaching primate characteristics, potentially reflecting greater reliance on visual inputs. Core hippocampal network dynamics—including proximo-distal gradients, pattern completion, and global remapping—remain largely conserved. This continuous trajectory provides a predictive framework for inferring human hippocampal coding from accessible model species. L, left eye monocular field; R, right eye monocular field.

| Fig. ID | Variable & Effect | Method | Statistics | P/Q value | Effect size | N number |
|---|---|---|---|---|---|---|
| Fig. 1d | Recording sites, TS vs. uniform sampling | Kolmogorov-Smirnov test | D = 0.114 | $P$ = 1.000 | - | 20 sites |
| Fig. 1d | Recording sites, RT vs. uniform sampling | Kolmogorov-Smirnov test | D = 0.089 | $P$ = 0.372 | - | 113 sites |
| Fig. 1g | LFP power, TS vs. RT. at 7 Hz | Wilcoxon rank-sum test | Z = −7.085 | $Q$ < 0.001 | $r$ = −0.429 | TS/RT: 149/124 experiments |
| Fig. 1g | LFP power, TS vs. RT. at 7.5 Hz | Wilcoxon rank-sum test | Z = −11.447 | $Q$ < 0.001 | $r$ = −0.693 | |
| Fig. 1g | LFP power, TS vs. RT. at 8 Hz | Wilcoxon rank-sum test | Z = −11.589 | $Q$ < 0.001 | $r$ = −0.701 | |
| Fig. 1g | LFP power, TS vs. RT. at 8.5 Hz | Wilcoxon rank-sum test | Z = −8.094 | $Q$ < 0.001 | $r$ = −0.490 | |
| Fig. 1g | LFP power, TS vs. RT. at 9 Hz | Wilcoxon rank-sum test | Z = −2.032 | $Q$ = 0.047 | $r$ = −0.123 | |
| Fig. 1g | LFP power, TS vs. RT. at 9.5 Hz | Wilcoxon rank-sum test | Z = 2.793 | $Q$ = 0.006 | $r$ = 0.169 | |
| Fig. 1g | LFP power, TS vs. RT. at 10 Hz | Wilcoxon rank-sum test | Z = 3.593 | $Q$ < 0.001 | $r$ = 0.217 | |
| Fig. 1j | Fraction of theta-locked neurons, TS vs. RT | chi-square test | $\chi^2(1) = 365.7$ | $P$ < 0.001 | φ = 0.323 | TS/RT: 1264/2248 cells |
| Fig. 1j | Fraction of theta-locked neurons, TS | chi-square test | $\chi^2(2) = 0.178$ | $P$ = 0.915 | - | 395/452/417 cells |
| Fig. 1j | Fraction of theta-locked neurons, RT | chi-square test | $\chi^2(2) = 1.542$ | $P$ = 0.463 | - | 735/806/707 cells |
| Fig. 1k | Theta phase-locking strength, TS vs. RT across subregions | Two-way RT-ANOVA | Species, $F(1, 1708) = 245.961$ | $P$ < 0.001 | $\eta^2 = 0.126$ | TS/RT: pCA1, 105/461 neurons |
| | | | Subregion, $F(2, 1708) = 2.723$ | $P$ = 0.066 | - | TS/RT: mCA1, 126/485 neurons |
| | | | Interaction, $F(2, 1708) = 2.168$ | $P$ = 0.115 | - | TS/RT: dCA1, 114/423 neurons |
| Fig. 2c | Linear speed (LS), TS vs. RT | Wilcoxon rank-sum test | Z = 13.328 | $P$ < 0.001 | $r$ = 0.614 | TS/RT: 293/178 experiments |
| Fig. 2d | Angular velocity (AV), TS vs. RT | Wilcoxon rank-sum test | Z = 12.166 | $P$ < 0.001 | $r$ = 0.561 | |
| Fig. 2e | Tortuosity, TS vs. RT | Wilcoxon rank-sum test | Z = −13.008 | $P$ < 0.001 | $r$ = −0.599 | |
| Fig. 2f | Thigmotaxis, TS vs. RT | Wilcoxon rank-sum test | Z = −6.813 | $P$ < 0.001 | $r$ = −0.314 | |
| Fig. 2g | AV vs. LS, TS vs. RT | Wilcoxon rank-sum test | Z = −9.912 | $P$ < 0.001 | $r$ = −0.457 | |
| Fig. 2h | Arena coverage, TS vs. RT | Wilcoxon rank-sum test | Z = −1.613 | $P$ = 0.107 | - | |
| Fig. 2i | Trajectory evenness, TS vs. RT | Wilcoxon rank-sum test | Z = 1.208 | $P$ = 0.227 | - | |
| Fig. 3c | Peak firing rate by cell, TS vs. RT across subregions | Two-way RT-ANOVA | Species, $F(1, 3895) = 635.509$ | $P$ < 0.001 | $\eta^2 = 0.140$ | TS/RT: pCA1, 422/863 neurons |
| | | | Subregion, $F(2, 3895) = 53.904$ | $P$ < 0.001 | $\eta^2 = 0.027$ | TS/RT: mCA1, 483/940 neurons |
| | | | Interaction, $F(2, 3895) = 9.625$ | $P$ < 0.001 | $\eta^2 = 0.005$ | TS/RT: dCA1, 420/773 neurons |
| Fig. 3c | Peak firing rate by recording site, TS vs. RT | Permutation test | - | $P$ = 0.296 | - | TS/RT: 20/113 sites |
| Fig. 3d | Spatial information content by cell, TS vs. RT across subregions | Two-way RT-ANOVA | Species, $F(1, 3895) = 2638.462$ | $P$ < 0.001 | $\eta^2 = 0.404$ | |
| | | | Subregion, $F(2, 3895) = 101.035$ | $P$ < 0.001 | $\eta^2 = 0.049$ | |
| | | | Interaction, $F(2, 3895) = 22.770$ | $P$ < 0.001 | $\eta^2 = 0.012$ | |
| Fig. 3d | SIc by recording site, TS vs. RT | Permutation test | - | $P$ = 0.109 | - | |

| Fig. ID | Variable & Effect | Method | Statistics | P/Q value | Effect size | N number |
|---|---|---|---|---|---|---|
| Fig. 3e | Coherence by cell, TS vs. RT across subregions | Two-way RT-ANOVA | Species, $F(1, 3895) = 1780.150$ | $P < 0.001$ | $\eta^2 = 0.314$ | TS/RT: pCA1, 422/863 neurons<br>TS/RT: mCA1, 483/940 neurons<br>TS/RT: dCA1, 420/773 neurons<br>TS/RT: 20/113 sites |
| | | | Subregion, $F(2, 3895) = 83.683$ | $P < 0.001$ | $\eta^2 = 0.041$ | |
| | | | Interaction, $F(2, 3895) = 5.480$ | $P = 0.004$ | $\eta^2 = 0.003$ | |
| Fig. 3e | Coherence by recording site, TS vs. RT | Permutation test | - | $P = 0.411$ | - | |
| Fig. 3f | Stability by cell, TS vs. RT across subregions | Two-way RT-ANOVA | Species, $F(1, 3887) = 1507.666$ | $P < 0.001$ | $\eta^2 = 0.280$ | |
| | | | Subregion, $F(2, 3887) = 84.963$ | $P < 0.001$ | $\eta^2 = 0.042$ | |
| | | | Interaction, $F(2, 3887) = 0.170$ | $P = 0.844$ | - | |
| Fig. 3f | Stability by recording site, TS vs. RT | Permutation test | - | $P = 0.173$ | - | |
| Fig. 3g | Fraction of place cells, TS vs. RT, pCA1 | chi-square test | $\chi^2(1) = 162.1$ | $P < 0.001$ | $\varphi = 0.355$ | TS/RT: 422/863 neurons |
| Fig. 3g | Fraction of place cells, TS vs. RT, mCA1 | chi-square test | $\chi^2(1) = 204.0$ | $P < 0.001$ | $\varphi = 0.379$ | TS/RT: 483/940 neurons |
| Fig. 3g | Fraction of place cells, TS vs. RT, dCA1 | chi-square test | $\chi^2(1) = 260.3$ | $P < 0.001$ | $\varphi = 0.467$ | TS/RT: 420/773 neurons |
| Fig. 3g | Fraction of place cells, TS subregions | chi-square test | $\chi^2(2) = 62.29$ | $P < 0.001$ | Cramer's $V = 0.217$ | 422/483/420 cells |
| Fig. 3g | Fraction of place cells, RT subregions | chi-square test | $\chi^2(2) = 33.87$ | $P < 0.001$ | Cramer's $V = 0.115$ | 863/940/773 cells |
| Fig. 4b | HD tuning by cell, TS vs. RT across subregions | Two-way RT-ANOVA | Species, $F(1, 3895) = 3.459$, | $P = 0.063$ | - | TS/RT: pCA1, 422/863 neurons<br>TS/RT: mCA1, 483/940 neurons<br>TS/RT: dCA1, 420/773 neurons<br>TS/RT: 20/113 sites |
| | | | Subregion, $F(2, 3895) = 145.172$ | $P < 0.001$ | $\eta^2 = 0.069$ | |
| | | | Interaction, $F(2, 3895) = 57.466$ | $P < 0.001$ | $\eta^2 = 0.029$ | |
| Fig. 4b | HD tuning by recording site, TS vs. RT | Permutation test | - | $P = 0.002$ | - | |
| Fig. 4c | Fraction of HD neurons, TS vs. RT, pCA1 | chi-square test | $\chi^2(1) = 245.0$ | $P < 0.001$ | $\varphi = 0.437$ | TS/RT: 422/863 neurons |
| Fig. 4c | Fraction of HD neurons, TS vs. RT, mCA1 | chi-square test | $\chi^2(1) = 213.8$ | $P < 0.001$ | $\varphi = 0.388$ | TS/RT: 483/940 neurons |
| Fig. 4c | Fraction of HD neurons, TS vs. RT, dCA1 | chi-square test | $\chi^2(1) =83.44$ | $P < 0.001$ | $\varphi = 0.265$ | TS/RT: 420/773 neurons |
| Fig. 4c | Fraction of HD neurons, TS subregions | chi-square test | $\chi^2(2) = 91.20$ | $P < 0.001$ | Cramer's $V = 0.262$ | 422/483/420 neurons |
| Fig. 4c | Fraction of HD neurons, RT subregions | chi-square test | $\chi^2(2) = 27.75$ | $P < 0.001$ | Cramer's $V = 0.104$ | 863/940/773 neurons |
| Main text | EB tuning vs. HD tuning, TS | Repeated-measures two-way RT-ANOVA | Metric, $F(1, 1322) = 104.519$ | $P < 0.001$ | $\eta^2 = 0.073$ | 422/483/420 neurons |
| | | | Subregion, $F(2, 1322) = 100.382$ | $P < 0.001$ | $\eta^2 = 0.132$ | |
| | | | Interaction, $F(2, 1322) = 32.266$ | $P < 0.001$ | $\eta^2 = 0.047$ | |
| Main text | EB tuning vs. HD tuning, RT | Repeated-measures two-way RT-ANOVA | Metric, $F(1, 2573) = 110.575$ | $P < 0.001$ | $\eta^2 = 0.041$ | 863/940/773 neurons |
| | | | Subregion, $F(2, 2573) = 17.044$ | $P < 0.001$ | $\eta^2 = 0.013$ | |
| | | | Interaction, $F(2, 2573) = 3.929$ | $P = 0.020$ | $\eta^2 = 0.003$ | |

| Fig. ID | Variable & Effect | Method | Statistics | P/Q value | Effect size | N number |
|---|---|---|---|---|---|---|
| | | | Species, $F(1, 3895) = 11.102$ | $P < 0.001$ | $\eta^2 = 0.003$ | TS/RT: pCA1, 422/863 neurons |
| Fig. 4d | EB tuning by cell, TS vs. RT across subregions | Two-way RT-ANOVA | Subregion, $F(2, 3895) = 51.846$ | $P < 0.001$ | $\eta^2 = 0.026$ | TS/RT: mCA1, 483/940 neurons |
| | | | Interaction, $F(2, 3895) = 12.364$ | $P < 0.001$ | $\eta^2 = 0.006$ | TS/RT: dCA1, 420/773 neurons |
| Fig. 4d | EB tuning by recording site, TS vs. RT | Permutation test | - | $P = 0.180$ | - | TS/RT: 20/113 sites |
| Fig. 4e | Fraction of EB neurons, TS vs. RT, pCA1 | chi-square test | $\chi^2(1) = 116.5$ | $P < 0.001$ | $\varphi = 0.301$ | TS/RT: 422/863 neurons |
| Fig. 4e | Fraction of EB neurons, TS vs. RT, mCA1 | chi-square test | $\chi^2(1) = 160.6$ | $P < 0.001$ | $\varphi = 0.336$ | TS/RT: 483/940 neurons |
| Fig. 4e | Fraction of EB neurons, TS vs. RT, dCA1 | chi-square test | $\chi^2(1) = 158.0$ | $P < 0.001$ | $\varphi = 0.364$ | TS/RT: 420/773 neurons |
| Fig. 4e | Fraction of EB neurons, TS subregions | chi-square test | $\chi^2(2) = 1.377$ | $P = 0.502$ | - | 422/483/420 cells |
| Fig. 4e | Fraction of EB neurons, RT subregions | chi-square test | $\chi^2(2) = 16.59$ | $P < 0.001$ | Cramer's $V$ = 0.080 | 863/940/773 cells |
| | | | Species, $F(1, 3895) = 218.330$ | $P < 0.001$ | $\eta^2 = 0.053$ | TS/RT: pCA1, 422/863 neurons |
| Fig. 4f | FL tuning by cell, TS vs. RT across subregions | Two-way RT-ANOVA | Subregion, $F(2, 3895) = 133.800$ | $P < 0.001$ | $\eta^2 = 0.064$ | TS/RT: mCA1, 483/940 neurons |
| | | | Interaction, $F(2, 3895) = 58.168$ | $P < 0.001$ | $\eta^2 = 0.029$ | TS/RT: dCA1, 420/773 neurons |
| Fig. 4f | FL tuning by recording site, TS vs. RT | Permutation test | - | $P = 0.006$ | - | TS/RT: 20/113 sites |
| Fig. 5c | Allocentric connections vs. others | Wilcoxon rank-sum test | Z = 2.237 | $P = 0.018$ | $r = 0.578$ | 15 combinations |
| Fig. 5f | Shuffle, TS vs. RT. at 60 s | Wilcoxon rank-sum test | Z = 3.972 | $Q = 0.001$ | $r = 0.172$ | |
| Fig. 5f | Shuffle, TS vs. RT. at 70 s | Wilcoxon rank-sum test | Z = 1.455 | $Q = 0.162$ | - | |
| Fig. 5f | Shuffle, TS vs. RT. at 80 s | Wilcoxon rank-sum test | Z = −0.615 | $Q = 0.539$ | - | |
| Fig. 5f | Shuffle, TS vs. RT. at 90 s | Wilcoxon rank-sum test | Z = −2.036 | $Q = 0.052$ | - | |
| Fig. 5f | Shuffle, TS vs. RT. at 100 s | Wilcoxon rank-sum test | Z = −2.995 | $Q = 0.004$ | $r = −0.130$ | TS/RT: 213/318 sessions |
| | | | Species, $F(1, 529) = 2.412$ | $P = 0.121$ | $\eta^2 = 0.005$ | |
| Fig. 5g | Decoding errors by session, TS vs. RT | Repeated-measures two-way RT-ANOVA | Alignment, $F(1, 529) = 1557.983$ | $P < 0.001$ | $\eta^2 = 0.747$ | |
| | | | Interaction, $F(1, 529) = 0.983$ | $P = 0.322$ | $\eta^2 = 0.002$ | |
| Fig. 5h | Neurons per decoding session, TS vs. RT | Wilcoxon rank-sum test | Z = −6.279 | $P < 0.001$ | $r = −0.272$ | |
| Fig. 5j | Neurons per session, TS vs. RT sub | Wilcoxon rank-sum test | $Z = 0.113$ | $P = 0.910$ | - | TS/RT: 213/213 sessions |
| Fig. 5j | Decoding accuracy, TS vs. RT subsampled | Wilcoxon rank-sum test | Z = −3.523 | $P < 0.001$ | $r = −0.171$ | |
| Fig. 6b | Hippocampal size, TS vs. RT | Welch's $t$-test | $t(3.545) = 3.378$ | $P = 0.034$ | Hedges' $g$ = 2.44 | TS/RT: 7/4 animals |
| Fig. 7b | AV across trials | Friedman test | $\chi^2(2) = 18.231$ | $P < 0.001$ | Kendall's $W$ = 0.351 | 26 experiments |
| Fig. 7c | Tortuosity across trials | Friedman test | $\chi^2(2) = 14.846$ | $P < 0.001$ | Kendall's $W$ = 0.286 | |

| Fig. ID | Variable & Effect | Method | Statistics | P/Q value | Effect size | N number |
|---|---|---|---|---|---|---|
| Fig. 7g | SIc, same vs.different conditions | Wilcoxon signed-rank test | Z = −1.049 | $P$ = 0.294 | - | 25 cells |
| Fig. 7g | Coherence, same vs.different conditions | Wilcoxon signed-rank test | Z = −1.293 | $P$ = 0.196 | - | |
| Fig. 7g | Stability, same vs.different conditions | Wilcoxon signed-rank test | Z = −1.323 | $P$ = 0.183 | - | |
| Fig. 7h | HD tuning, same vs.different conditions | Wilcoxon signed-rank test | Z = −0.436 | $P$ = 0.671 | - | 19 cells |
| Fig. 7i | FL tuning, same vs.different conditions | Wilcoxon signed-rank test | Z = 2.170 | $P$ = 0.023 | $r$ = 0.686 | 10 cells |
| Fig. 8a | Fraction of active neurons, TS vs. RT | chi-square test | $\chi^2(1) = 3.542$ | $P$ = 0.060 | - | TS/RT: 188/383 cells |
| Fig. 8d | Room-switch ΔSC by SIc group, TS vs. RT | Two-way RT-ANOVA | Species, $F(1, 958) = 17.200$<br>Group, $F(2, 958) = 74.524$<br>Interaction, $F(2, 958) = 1.103$ | $P < 0.001$<br>$P < 0.001$<br>$P$ = 0.332 | $\eta^2 = 0.018$<br>$\eta^2 = 0.135$<br>- | non-positional, 155/34 neurons low-positional, 148/91 neurons high-positional, 79/492 neurons |
| Fig. 8g | Dirction distribution by cell, TS176 | Rayleigh test | Z = 4.112 | $P$ = 0.011 | - | 7 neurons |
| Fig. 8g | Dirction distribution by cell, TS177 | Rayleigh test | Z = 6.507 | $P < 0.001$ | - | 18 neurons |
| Fig. 8g | Dirction distribution by cell, TS180 | Rayleigh test | Z = 1.868 | $P$ = 0.156 | - | 6 neurons |
| Fig. 8g | Dirction distribution by cell, TS181 | Rayleigh test | Z = 19.609 | $P < 0.001$ | - | 54 neurons |
| Fig. 8g | Dirction distribution by cell, TS209 | Rayleigh test | Z = 4.125 | $P$ = 0.010 | - | 7 neurons |
| Fig. 8g | Dirction distribution by cell, TS224 | Rayleigh test | Z = 9.788 | $P < 0.001$ | - | 12 neurons |
| Fig. 8g | Dirction distribution by cell, TS230 | Rayleigh test | Z = 2.930 | $P$ = 0.037 | - | 3 neurons |
| Fig. 8g | Dirction distribution by cell, TS238 | Rayleigh test | Z = 0.959 | $P$ = 0.396 | - | 8 neurons |
| Fig. 8h | Dirction distribution by cell, all cells | Rayleigh test | Z = 47.317 | $P < 0.001$ | - | 115 neurons |
| Fig. 8i | Pairwise differences by experiments | Moore's paired-sample circular test | R* = 0.773 | $P$ = 0.181 | - | 30 experiments |

**Supplementary Table 1 | Statistical analyses and outcomes**

| Fig. ID | Variable & Effect | Two-way RT-ANOVA | | Linear Mixed-effects Model | | |
|---|---|---|---|---|---|---|
| | | Statistics | *P* value | Statistics | *P* value | Animal effect |
| Fig. 1 | **Theta-phase-locking mean vector length** | | | | | |
| | Species | F (1, 1708) = 245.961 | *P* < 0.001 | F (1, 22.477) = 59.239 | *P* < 0.001 | *P* = 0.034 |
| | Subregion | F (2, 1708) = 2.723 | *P* = 0.066 | F (2, 70.160) = 1.085 | *P* = 0.344 | |
| | Species × Subregion | F (2, 1708) = 2.168 | *P* = 0.115 | F (2, 70.160) = 0.465 | *P* = 0.630 | |
| Fig. 3 | **Peak firing rate** | | | | | |
| | Species | F (1, 3895) = 635.509 | *P* < 0.001 | F (1, 25.332) = 18.724 | *P* < 0.001 | *P* = 0.001 |
| | Subregion | F (2, 3895) = 53.904 | *P* < 0.001 | F (2, 54.280) = 1.582 | *P* = 0.215 | |
| | Species × Subregion | F (2, 3895) = 9.625 | *P* < 0.001 | F (2, 54.280) = 1.438 | *P* = 0.246 | |
| | **Spatial information content** | | | | | |
| | Species | F (1, 3895) = 2638.462 | *P* < 0.001 | F (1, 27.591) = 165.788 | *P* < 0.001 | *P* = 0.001 |
| | Subregion | F (2, 3895) = 101.035 | *P* < 0.001 | F (2, 63.549) = 11.038 | *P* < 0.001 | |
| | Species × Subregion | F (2, 3895) = 22.770 | *P* < 0.001 | F (2, 63.549) = 3.315 | *P* = 0.043 | |
| | **Spatial coherence** | | | | | |
| | Species | F (1, 3895) = 1780.150 | *P* < 0.001 | F (1, 28.928) = 68.794 | *P* < 0.001 | *P* = 0.001 |
| | Subregion | F (2, 3895) = 83.683 | *P* < 0.001 | F (2, 61.925) = 4.028 | *P* = 0.023 | |
| | Species × Subregion | F (2, 3895) = 5.480 | *P* = 0.004 | F (2, 61.925) = 3.189 | *P* = 0.048 | |
| | **Spatial stability** | | | | | |
| | Species | F (1, 3887) = 1507.666 | *P* < 0.001 | F (1, 29.464) = 74.244 | *P* < 0.001 | *P* = 0.001 |
| | Subregion | F (2, 3887) = 84.963 | *P* < 0.001 | F (2, 64.031) = 6.261 | *P* = 0.003 | |
| | Species × Subregion | F (2, 3887) = 0.170 | *P* = 0.844 | F (2, 64.031) = 2.764 | *P* = 0.071 | |
| Fig. 4 | **Head direction mean vector length** | | | | | |
| | Species | F (1, 3895) = 3.459 | *P* = 0.063 | F (1, 29.714) = 0.066 | *P* = 0.799 | *P* < 0.001 |
| | Subregion | F (2, 3895) = 145.172 | *P* < 0.001 | F (2, 63.834) = 7.619 | *P* = 0.001 | |
| | Species × Subregion | F (2, 3895) = 57.466 | *P* < 0.001 | F (2, 63.834) = 7.468 | *P* = 0.001 | |
| | **Egocentric bearing mean vector length** | | | | | |
| | Species | F (1, 3895) = 11.102 | *P* < 0.001 | F (1, 27.316) = 1.021 | *P* = 0.321 | *P* = 0.002 |
| | Subregion | F (2, 3895) = 51.846 | *P* < 0.001 | F (2, 65.612) = 4.003 | *P* = 0.023 | |
| | Species × Subregion | F (2, 3895) = 12.364 | *P* < 0.001 | F (2, 65.612) = 3.443 | *P* = 0.038 | |
| | **Facing location mean vector length** | | | | | |
| | Species | F (1, 3895) = 218.330 | *P* < 0.001 | F (1, 30.038) = 17.527 | *P* < 0.001 | *P* = 0.001 |
| | Subregion | F (2, 3895) = 133.800 | *P* < 0.001 | F (2, 68.066) = 10.961 | *P* < 0.001 | |
| | Species × Subregion | F (2, 3895) = 58.168 | *P* < 0.001 | F (2, 68.066) = 12.321 | *P* < 0.001 | |
| Extended Data Fig. 3 | **Mean firing rate** | | | | | |
| | Species | F (1, 3895) = 35.547 | *P* < 0.001 | F (1, 28.920) = 6.532 | *P* = 0.016 | *P* = 0.001 |
| | Subregion | F (2, 3895) = 24.081 | *P* < 0.001 | F (2, 66.758) = 4.492 | *P* = 0.015 | |
| | Species × Subregion | F (2, 3895) = 30.327 | *P* < 0.001 | F (2, 66.758) = 3.465 | *P* = 0.037 | |
| | **Receptive field size** | | | | | |
| | Species | F (1, 3895) =143.664 | *P* < 0.001 | F (1, 29.860) = 17.651 | *P* < 0.001 | *P* = 0.002 |
| | Subregion | F (2, 3895) = 51.818 | *P* < 0.001 | F (2, 73.617) = 11.075 | *P* < 0.001 | |
| | Species × Subregion | F (2, 3895) = 21.608 | *P* < 0.001 | F (2, 73.617) = 2.208 | *P* = 0.117 | |

| Fig. ID | Variable & Effect | Two-way RT-ANOVA Statistics | Two-way RT-ANOVA *P* value | Linear Mixed-effects Model Statistics | Linear Mixed-effects Model *P* value | Animal effect |
|---|---|---|---|---|---|---|
| Extended Data Fig. 3 | **Receptive field number** | | | | | |
| | Species | F (1, 3895) = 82.687 | *P* < 0.001 | F (1, 24.099) = 4.557 | *P* = 0.043 | *P* = 0.006 |
| | Subregion | F (2, 3895) = 11.121 | *P* < 0.001 | F (2, 59.741) = 0.838 | *P* = 0.438 | |
| | Species × Subregion | F (2, 3895) = 17.159 | *P* < 0.001 | F (2, 59.741) = 5.851 | *P* = 0.005 | |
| | **Spatial information rate** | | | | | |
| | Species | F (1, 3895) = 1362.754 | *P* < 0.001 | F (1, 25.877) = 42.800 | *P* < 0.001 | *P* = 0.001 |
| | Subregion | F (2, 3895) = 83.272 | *P* < 0.001 | F (2, 55.127) = 3.573 | *P* = 0.035 | |
| | Species × Subregion | F (2, 3895) = 14.198 | *P* < 0.001 | F (2, 55.127) = 2.676 | *P* = 0.078 | |
| Extended Data Fig. 4 | **Mean firing rate (position-selective neurons)** | | | | | |
| | Species | F (1, 3142) = 55.626 | *P* < 0.001 | F (1, 29.956) = 10.427 | *P* = 0.003 | *P* = 0.003 |
| | Subregion | F (2, 3142) = 18.387 | *P* < 0.001 | F (2, 71.794) = 6.496 | *P* = 0.003 | |
| | Species × Subregion | F (2, 3142) = 20.218 | *P* < 0.001 | F (2, 71.794) = 6.957 | *P* = 0.002 | |
| | **Peak firing rate (position-selective neurons)** | | | | | |
| | Species | F (1, 3142) = 291.523 | *P* < 0.001 | F (1, 27.652) = 13.424 | *P* = 0.001 | *P* = 0.002 |
| | Subregion | F (2, 3142) = 35.009 | *P* < 0.001 | F (2, 61.378) = 3.090 | *P* = 0.053 | |
| | Species × Subregion | F (2, 3142) = 9.753 | *P* < 0.001 | F (2, 61.378) = 2.192 | *P* = 0.120 | |
| | **Place field size (position-selective neurons)** | | | | | |
| | Species | F (1, 3142) = 193.714 | *P* < 0.001 | F (1, 25.866) = 33.837 | *P* < 0.001 | *P* = 0.013 |
| | Subregion | F (2, 3142) = 23.782 | *P* < 0.001 | F (2, 66.950) = 10.344 | *P* < 0.001 | |
| | Species × Subregion | F (2, 3142) = 19.346 | *P* < 0.001 | F (2, 66.950) = 6.468 | *P* = 0.003 | |
| | **Place field number (position-selective neurons)** | | | | | |
| | Species | F (1, 3142) = 39.615 | *P* < 0.001 | F (1, 21.981) = 7.593 | *P* = 0.012 | *P* = 0.040 |
| | Subregion | F (2, 3142) = 3.809 | *P* = 0.022 | F (2, 59.605) = 0.913 | *P* = 0.407 | |
| | Species × Subregion | F (2, 3142) = 7.831 | *P* < 0.001 | F (2, 59.605) = 4.578 | *P* = 0.014 | |
| | **Spatial information content (position-selective neurons)** | | | | | |
| | Species | F (1, 3142) = 1486.849 | *P* < 0.001 | F (1, 28.898) = 149.032 | *P* < 0.001 | *P* = 0.004 |
| | Subregion | F (2, 3142) = 37.570 | *P* < 0.001 | F (2, 70.374) = 10.523 | *P* < 0.001 | |
| | Species × Subregion | F (2, 3142) = 15.281 | *P* < 0.001 | F (2, 70.374) = 4.281 | *P* = 0.018 | |
| | **Spatial information rate (position-selective neurons)** | | | | | |
| | Species | F (1, 3142) = 664.543 | *P* < 0.001 | F (1, 26.510) = 32.343 | *P* < 0.001 | *P* = 0.002 |
| | Subregion | F (2, 3142) = 43.456 | *P* < 0.001 | F (2, 57.995) = 5.183 | *P* = 0.008 | |
| | Species × Subregion | F (2, 3142) = 12.438 | *P* < 0.001 | F (2, 57.995) = 3.874 | *P* = 0.026 | |
| | **Spatial coherence (position-selective neurons)** | | | | | |
| | Species | F (1, 3142) = 905.980 | *P* < 0.001 | F (1, 31.465) = 54.886 | *P* < 0.001 | *P* = 0.001 |
| | Subregion | F (2, 3142) = 24.105 | *P* < 0.001 | F (2, 69.572) = 3.084 | *P* = 0.052 | |
| | Species × Subregion | F (2, 3142) = 4.123 | *P* = 0.016 | F (2, 69.572) = 2.819 | *P* = 0.067 | |
| | **Spatial stability (position-selective neurons)** | | | | | |
| | Species | F (1, 3138) = 678.055 | *P* < 0.001 | F (1, 33.161) = 57.116 | *P* < 0.001 | *P* = 0.001 |
| | Subregion | F (2, 3138) = 26.300 | *P* < 0.001 | F (2, 75.121) = 4.424 | *P* = 0.015 | |
| | Species × Subregion | F (2, 3138) = 1.340 | *P* = 0.262 | F (2, 75.121) = 2.905 | *P* = 0.061 | |

| Fig. ID | Variable & Effect | Two-way RT-ANOVA | | Linear Mixed-effects Model | | |
|---|---|---|---|---|---|---|
| | | Statistics | $P$ value | Statistics | $P$ value | Animal effect |
| Extended Data Fig. 4 | **HD Mean vector length (HD-selective neurons)** | | | | | |
| | Species | F (1, 1371) = 1.793 | $P = 0.181$ | F (1, 27.775) = 0.698 | $P = 0.410$ | $P = 0.004$ |
| | Subregion | F (2, 1371) = 32.870 | $P < 0.001$ | F (2, 65.547) = 4.175 | $P = 0.020$ | |
| | Species × Subregion | F (2, 1371) = 9.229 | $P < 0.001$ | F (2, 65.547) = 3.280 | $P = 0.044$ | |
| | **EB Mean vector length (EB-selective neurons)** | | | | | |
| | Species | F (1, 995) = 7.460 | $P = 0.006$ | F (1, 23.198) = 1.133 | $P = 0.298$ | $P = 0.012$ |
| | Subregion | F (2, 995) = 15.327 | $P < 0.001$ | F (2, 59.491) = 2.526 | $P = 0.089$ | |
| | Species × Subregion | F (2, 995) = 8.660 | $P < 0.001$ | F (2, 59.491) = 3.089 | $P = 0.053$ | |
| | **FL Mean vector length (FL-selective neurons)** | | | | | |
| | Species | F (1, 2170) = 183.156 | $P < 0.001$ | F (1, 27.894) = 13.726 | $P = 0.001$ | $P = 0.005$ |
| | Subregion | F (2, 2170) = 36.444 | $P < 0.001$ | F (2, 66.758) = 7.726 | $P = 0.001$ | |
| | Species × Subregion | F (2, 2170) = 28.195 | $P < 0.001$ | F (2, 66.758) = 4.731 | $P = 0.012$ | |
| Extended Data Fig. 5 | **Pos log-likelihood increase** | | | | | |
| | Species | F (1, 2106) = 854.948 | $P < 0.001$ | F (1, 35.216) = 108.611 | $P < 0.001$ | $P = 0.003$ |
| | Subregion | F (2, 2106) = 34.249 | $P < 0.001$ | F (2, 84.409) = 9.757 | $P < 0.001$ | |
| | Species × Subregion | F (2, 2106) = 0.859 | $P = 0.424$ | F (2, 84.409) = 0.586 | $P = 0.559$ | |
| | **HD log-likelihood increase** | | | | | |
| | Species | F (1, 2106) = 1593.863 | $P < 0.001$ | F (1, 30.479) = 134.188 | $P < 0.001$ | $P = 0.004$ |
| | Subregion | F (2, 2106) = 6.545 | $P = 0.001$ | F (2, 71.793) = 2.447 | $P = 0.094$ | |
| | Species × Subregion | F (2, 2106) = 1.150 | $P = 0.317$ | F (2, 71.793) = 1.866 | $P = 0.162$ | |
| | **FL log-likelihood increase** | | | | | |
| | Species | F (1, 2106) = 650.552 | $P < 0.001$ | F (1, 31.876) = 86.114 | $P < 0.001$ | $P = 0.007$ |
| | Subregion | F (2, 2106) = 11.959 | $P < 0.001$ | F (2, 79.172) = 4.863 | $P = 0.010$ | |
| | Species × Subregion | F (2, 2106) = 1.188 | $P = 0.305$ | F (2, 79.172) = 2.729 | $P = 0.071$ | |
| | **EB log-likelihood increase** | | | | | |
| | Species | F (1, 2106) = 1237.615 | $P < 0.001$ | F (1, 30.862) = 223.989 | $P < 0.001$ | $P = 0.014$ |
| | Subregion | F (2, 2106) = 1.279 | $P = 0.278$ | F (2, 78.814) = 2.489 | $P = 0.089$ | |
| | Species × Subregion | F (2, 2106) = 16.817 | $P < 0.001$ | F (2, 78.814) = 3.937 | $P = 0.023$ | |

**Supplementary Table 2 | Comparison of two-way RT-ANOVA and linear mixed-effects models.**

Both methods show similar species effects; subregion effects are attenuated in some mixed-effects comparisons, possibly due to over-correction for inter-animal recording site variation.

**Supplementary Video 1 | Tree shrew open-field navigation.**

Grayscale camera recording of a tree shrew navigating while chasing randomly distributed rewards, synchronized with electrophysiology and tracking experiments.
